# Consistent estimation of complete neuronal connectivity in large neuronal populations using sparse “shotgun” neuronal activity sampling

**DOI:** 10.1101/032409

**Authors:** Yuriy Mishchenko

## Abstract

We investigate the properties of the recently proposed “shotgun” sampling approach for the common inputs problem in the functional estimation of neuronal connectivity. We study the asymptotic correctness, the speed of convergence, and the data size requirements of such an approach. We find that the shotgun approach can be expected to allow the inference of the complete connectivity matrix in large neuronal populations under some rather general conditions. However, we find that the posterior error of the shotgun connectivity estimator may grow quickly with the size of the unobserved neuronal populations, the connectivity strength, and the square of the observations’ sparseness. This implies that the shotgun connectivity estimation will require significant amounts of neuronal activity data whenever the number of neurons in the observed populations is small. We present a numerical approach for solving the shotgun estimation problem in general settings and use it to demonstrate the shotgun connectivity inference in simulated synfire and weakly coupled cortical neuronal networks.

## 1 Introduction

Recent advances in multi-neuronal activity recordings using calcium-sensitive fluorescence imaging have made it possible to image the activity of large neuronal populations over extended periods of time (Tsien, 1989; Yuste et al., 2006; Cossart et al., 2003; Ohki et al., 2005; Reddy et al., 2008a; Grewe et al., 2010). Bulk-loading of organic calcium-sensitive fluorescent dyes offers the fluorescent signal-to-noise ratio (SNR) sufficient for resolving individual action potentials (spikes) of neurons (Yuste et al., 2006; Stosiek et al., 2003) and genetically encoded calcium indicators are approaching the SNR levels necessary for neuronal activity imaging with single spike accuracy (Wallace et al., 2008). Modern cooled CCD-microscopy can allow imaging of calcium fluorescence in neuronal populations in-vitro with frame-rates of over 60 Hz (Djurisic et al., 2004) and 2-photon laser scanning microscopy offers similar frame-rates in-vivo (Iyer et al., 2006; Salome et al., 2006; Reddy et al., 2008b; Cotton et al., 2013; Theis et al., 2015). These advances now allow studying the single-cell structure of neuronal circuits in the brain using accurate statistical approaches (Pillow et al., 2008; Stevenson et al., 2008a; Stevenson et al., 2008b; Stevenson et al., 2009; Mishchenko et al., 2011).

One of the biggest challenges of the functional analysis of the neuronal connectivity in he brain remains the presence of unobserved or hidden inputs in the recordings of neuronal population activity (Nykamp, 2007; Pillow and Latham, 2007; Vidne et al., 2009). Because functional connectivity estimation relies on correlating the activity of different neurons in a neuronal population over extended periods of time, the presence of unobserved inputs can contribute errors to functional analysis. In particular, the well-known “common inputs” problem causes one to mistake the correlations in the activity of different neurons caused by the correlations in their unobserved inputs for a direct connection between those neurons (Nykamp, 2005; Nykamp, 2007; Kulkarni and Paninski, 2007; Pillow and Latham, 2007). Despite rapid progress in experimental neuronal population activity imaging techniques, the simultaneous observation of the activity of all neurons even in the smallest of neuronal circuits is currently not plausible, and the development of robust analytical and computational techniques for overcoming the hidden inputs problem remains one of the important open questions of computational neuroscience (Nykamp, 2005; Nykamp, 2007; Kulkarni and Paninski, 2007; Pillow and Latham, 2007; Vidne et al., 2009; Keshri et al., 2013).

Recently a promising approach for overcoming the hidden inputs problem—the shotgun sampling—had been proposed in (Turaga et al., 2013; Keshri et al., 2013). In this approach, neurons in a large neuronal population are proposed to be imaged in small random groups, whereas the connectivity matrix of the complete neuronal population is assembled statistically by combining the information about the neuronal connectivity from different such partial measurements. The shotgun approach offers the possibility for reconstructing the connectivity of large neuronal circuits by using limited imaging resources, without the need to simultaneously image the entire neuronal circuits.

In this paper, we perform a systematic analysis of certain aspects of the shotgun sampling proposal such as the asymptotic correctness of the connectivity estimation, the speed of convergence, and the necessary data sizes. It may not be clear at first if the shotgun connectivity estimation is really free from the hidden inputs problem. One may observe that, even as all neurons in a neuronal population do get imaged with this approach over different points of time, these observations still fail to provide the complete input-output relationships of even a single neuron in the population: the total number of neurons that need to be simultaneously imaged to provide all the inputs of even a single neuron in mammalian cortex can be as high as 10,000. Without having the information about the complete set of inputs and outputs of any single neuron, one may wonder if the hidden inputs problem is really resolved. Furthermore, if the shotgun approach does allow the unambiguous determination of the complete connectivity matrix, what are the trade-offs that had been made? In particular, what is the minimal imaging time required for a given accuracy of the connectivity matrix reconstruction and how does this time scale with the size of the unobserved populations, observation sparseness, and other parameters?

Here, we show that the shotgun approach can be expected to recover the complete neuronal connectivity matrix in general neuronal activity models under some rather general conditions. We calculate the speed of convergence of the shotgun connectivity estimator and show that the imaging time required to achieve a given estimation accuracy scales with the number of neurons in the unobserved neuronal populations as well as the inverse square of the fraction of neurons observed during one imaging trial. This scaling is inopportune for the reconstructions of neuronal connectivity in the situations, where the number of unobserved neurons remains large. We further discuss a numerical approach for solving the connectivity estimation problem from shotgun sampling data and use it to demonstrate the shotgun connectivity estimation in small model neuronal populations.

In this paper, we focus specifically on the problem of the inference of the underlying neuronal connectivity matrix from partial neuronal population activity observations. In the case of calcium imaging—currently the most plausible modality for collecting large scale neuronal population activity data—another important component of such inference is the deconvolution of the calcium signal into the underlying neuronal spike trains. In recent years, significant progress had been made with respect to the deconvolution of the calcium imaging signal, for example, see (Vogelstein et al., 2009) and (Vogelstein et al., 2010), and (Theis et al., 2015) for a survey and comparison of recent deconvolution approaches. Unfortunately, recent experimental work had indicated that the existing calcium signal deconvolution approaches still may need significant improvement to be practically usable (Cotton et al., 2013; Theis et al., 2015). In this paper, we do not consider this important aspect of the analysis of functional data. Instead, we focus on the mathematical and computational problem of specifically the shotgun estimation, assuming that the deconvolved neuronal activity data is already available (Section 2.4) or that an effective neuronal activity model based on the continuous calcium fluorescence signal can be used (Section 2.2). However, in the future the incorporation of the calcium imaging deconvolution problem into and the impact of this problem onto the inference of neuronal connectivity will require more detailed investigation, including the understanding of the impact of the added uncertainties in the spike timing (as deduced from a lower frame-rate calcium imaging data), the apparently high fractions of lost spikes and the tendency to overestimate bursting and underestimate isolated neuronal spiking activity (Cotton et al., 2013; Theis et al., 2015).

In the remainder of the paper our presentation is organized as follows. In Materials and Methods, Section 2.1, we provide a general overview of the shotgun neuronal connectivity estimation problem and offer its mathematical formalization. In Section 2.2, we investigate the shotgun connectivity estimation problem in a linear neuronal population activity model. We analytically obtain the properly marginalized observations log-likelihood in this model in asymptotic limit and derive the conditions under which the maximum likelihood estimator in this model is consistent. Furthermore, we explicitly derive the sufficient conditions for the consistency of the shotgun connectivity estimation in general spiking models of neuronal activity, where the rate of neuronal spiking is described as a general function of linearly summed inputs. We also analytically demonstrate and discuss the impact of the hidden inputs problem on the connectivity estimation in the linear model. In Section 2.3, we study the shotgun connectivity estimation in general settings and propose an exact numerical algorithm for solving the associated estimation problem in general causal neuronal population activity models. In Section 2.4, we discuss the numerical simulations performed in this paper.

In Results, Section 3.1, we generalize the results of Section 2.2 to show that the shotgun maximum likelihood estimator can be proved more generally to converge to the true connectivity matrix of a neuronal population as long as certain conditions are met by the set of the neuronal subpopulations imaged by the shotgun sampling. In Section 3.2, we use this result to propose a different organization of the shotgun neuronal activity sampling that may be more advantageous for experimental realization. In Sections 3.3 and 3.4, we discuss the results of the shotgun connectivity estimation in simulated linear and generalized linear neuronal models. Discussion and conclusions follow in Section 4.

## 2 Materials and Methods

### 2.1 The shotgun sampling approach for neuronal connectivity estimation

In the shotgun sampling, one’s objective is to recover the effective connectivity matrix of a large neuronal population using a collection of partial samples of the activity of that population’s different subpopulations. One simple way to visualize this idea is to think about the reconstruction of a large image by using small fragments of that image from the image’s different locations.

More formally, we define the effective connectivity matrix 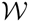 as a parameter of a statistical model of neuronal population activity, 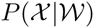, where 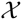 stands for the raster of the historical activities of all neurons in a neuronal population, and 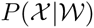 is the model likelihood of observing a particular raster 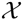. In network models, 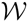 is typically a matrix 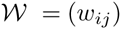 having as many rows and columns as there are neurons in the population, *N*. Each element *w_ij_* characterizes the impact of the past activity of one “input” neuron *j* on the activity of one “output” neuron *i*. Our objective here, therefore, is to estimate the complete matrix 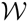 given a series of partial observations of the activity 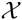.

In the context of this problem we introduce the following notation. We denote the raster of the activities of the entire neuronal population (that is, both observed and hidden) over all observations by the subscript-less symbol 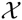, and we denote the activity of that population during one observation *t* by the symbol 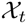. That is, 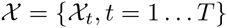, where *T* is the number of the observations. Similarly, we denote the collection of all observed neuronal activities over all observations by the subscript-less symbol *X*, whereas the activity of the observed neurons during one shotgun observation *t* is denoted by the symbol *X_t_*, so that *X* = {*X_t_, t* = 1 … *T*}. Finally, the collection of all unobserved neuronal activities will be referred to by using the symbols *Y* and *Y_t_*, respectively, so that *Y* = {*Y_t_, t* = 1 …*T*}.

We formally say that the shotgun estimation problem is the problem of estimating the effective connectivity matrix 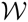 of a statistical model of neuronal population activity 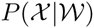 given a set of partial observations of neuronal activity *X*.

For the simplicity of the following discussion, in a significant part of this manuscript we will focus on the shotgun connectivity estimation formulated for a single output neuron and a randomly sampled population of neuronal inputs. In this picture, the output neuron is observed continuously in every observation, whereas the set of observed neuronal inputs changes. This assumption allows us to significant simplify the discussion, while not leading to any significant loss of generality. The latter is because in typical network models of neuronal activity the activities of neurons are conditionally independent given the activity of the presynaptic neuronal population and the respective input connection weights. In other words, the full likelihood 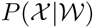 in such models factorizes over the rows of the connectivity matrix 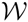,

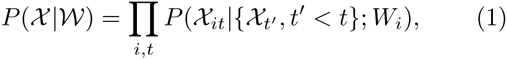
where *W_i_* = {*w_ij_*, *j* = 1 … *N*}. It is possible, therefore, to perform the estimation of 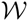 one row *W_i_* at a time.

In the rest of the paper we will consistently make use of the following notation. The script symbol 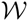 will always refer to the complete connectivity matrix of the entire neuronal population of *N* neurons, whereas the symbols *W_i_* will refer to a single row of 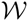, corresponding to the set of all input connection weights of one neuron. The symbol *w_ij_* will refer to one connection weight between an output neuron *i* and an input neuron *j*.

The script symbols 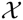 and 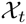 will be always used to refer to the neuronal activity of the entire neuronal population, whereas the plain symbols *X, X_t_, Y*, and *Y_t_* will be used to denote the observed and the unobserved parts of the neuronal activity, respectively. We shall occasionally make use of the symbols *X_t_* and *Y_t_* to refer to the set of neurons contained in *X_t_* or *Y_t_*, as opposed to their activities. This distinction will always be made clear by the context. Finally, 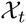, *X_t_* and *Y_t_* will be understood to be column-vectors, while *W_i_* will always be a row-vector.

### 2.2 The shotgun connectivity estimation in linear neuronal population activity model

#### 2.2.1 The linear model of neuronal activity

The linear model of neuronal activity has the advantage here of allowing the analysis of the shotgun connectivity estimation approach to be carried out analytically. The primary reason warranting the inspection of this model here, thus, is its analytical tractability.

In the linear neuronal activity model, we model the activity of a neuronal population using a continuous variable 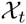, which can be thought, for example, to represent suitably smoothed firing rates of different neurons or the standard Δ*F/F* calcium imaging measure of neuronal activity. The input-output relationships between these variables is assumed to be linear,

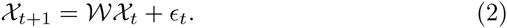

Thus, here 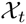 is an *N*-element column-vector representing the activity of the population of *N* neurons, while the index *t* plays the role of discrete time. The parameter 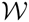 is a *N* × *N* connectivity matrix, and *ϵ* is a i.i.d. normal random noise variable. Without loss of generality, we assume *ϵ* to have the variances of one. The model probability density 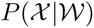 then would be defined by,

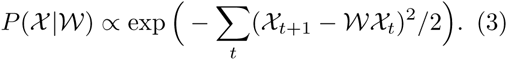

The problem of estimating the connectivity matrix 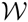 is simple, if the activity of the entire neuronal population is observed. In that case, we consider the observations’ log-likelihood,

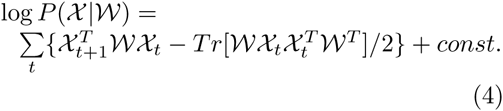

The maximum likelihood estimation (MLE) of 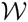 is found as the maximum of Eq. (4). One can see rather immediately that such MLE converges to the true connectivity matrix whenever the matrix 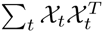 is of full rank.

In the partial observations case, a similar program can be pursued but one has to properly consider now the marginalized likelihood 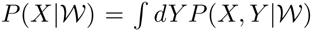, where the unobserved neural activity variables *Y* had been integrated out. For the linear neuronal activity model, we can calculate this likelihood analytically (that is, in the asymptotic limit of large numbers of observations) and study the properties of such MLE explicitly.

#### 2.2.2 The calculation of the partial observations likelihood in the linear neuronal activity model

In this section, we aim to calculate the marginalized observations likelihood 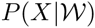 for the shotgun connectivity estimation in the linear neuronal activity model. We specifically focus on the input-output form of the model (2), with a scalar output variable *Z_it_* and two vector input variables *X_t_* and *Y_t_*, defined by the relationship,

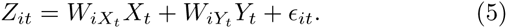

Here, *Z_it_* represents the “output” activity of one neuron *i* at time *t* + 1, while *X_t_* with *Y_t_* represent the activities of the observed and the unobserved “input” neuronal populations at time *t*. *W_i_* is the row of the complete connectivity matrix 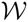 specifying the input connection weights of the output neuron *i*, and 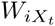 and 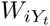 are the parts of *W_i_* corresponding to the observed and the unobserved inputs *X_t_* and *Y_t_*, respectively. *ϵ_it_* is an i.i.d. zero mean and unit variance normal random variable representing noise.

The inputs *X_t_* and *Y_t_* are assumed to be drawn jointly from a distribution *P*(*X_t_, Y_t_*). Understanding that *X_t_* and *Y_t_* represent the activity of the neuronal population in the model given by Eq. (2), the distribution *P*(*X_t_, Y_t_*) can be taken as the stationary distribution of the Markov process defined by Eq. (2). This distribution is Gaussian. By suitably offsetting the variables 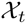 in Eq.(2), it is always possible to make this distribution have vanishing first moments. However, the covariance matrix of this disribution, *Σ*, in general will be nontrivial.

It shall be noted that in the above the connectivity vector *W_i_* and the covariance matrix *Σ* are both the parameters of the model (5). However, it can be also observed that *Σ* is not an independent parameter per-se since, by virtue of *P*(*X_t_, Y_t_*) being the stationary distribution of (2), it depends on the connectivity matrix 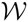. We will ignore this point for now, assuming no connection between 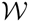 and *Σ*, as this will have no impact on the complexity of the calculations that we will need to perform, while a more general result can be obtained by ignoring this connection. However, see Section 2.2.5 for some important repercussions.

The specific quantity of interest in this section is the average log-likelihood of the observed data *Z_i_* and *X* in model (5) (in the sense of the compositional estimation of 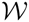 one output neuron at a time, see Section 2.1), marginal over the missing data represented by the hidden input variables *Y*,

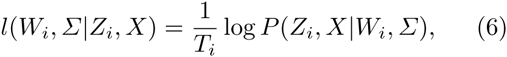
where *P*(*Z_i_, X|W_i_, Σ*) is the marginal distribution of the observed variables in model (5), *P*(*Z_i_, X|W_i_, Σ*) *= ∫ dY P*(*Z_i_, X, Y|W_i_, Σ*). Note that *Z_i_* and *X* in Eq. (6) are the collections of all observations (*Z_it_*, *X_t_*) for all 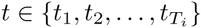 such that the output neuron *i* is observed, per the general conventions of Section 2.1. Therefore, in the RHS of Eq. (6) *Z_i_, X*, and *P*(*Z_i_*, *X*| *W_i_, Σ*) are all dependent on *T_i_* in this manner. Also note that both *Z_i_* and *X* are the observed data.

When the number of observations *T_i_* is large, the RHS of Eq. (6) converges in probability to the expectation value of log *P*(*Z_it_*, *X_t_|W_i_*, *Σ*) under the true distribution of the observed inputs and outputs *P*(*Z_it_*, *X_t_*),

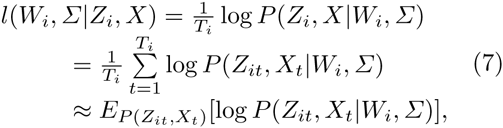
in which we recognize the expected log-likelihood function,

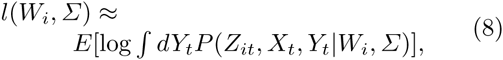
where the expectation, again, is taken with respect to the true density of the observed input and output variables, *X_t_* and *Z_it_*, and we used

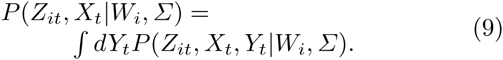

We see that it is necessary for us now to calculate,

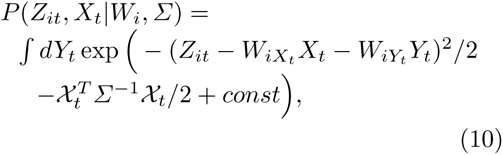
where 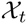 is the vector of the complete input activities formed by suitably combining *X_t_* and *Y_t_*, that is, 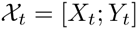. The integral in Eq. (10) can be taken explicitly, although the respective calculation is complex and we move it to Appendix B. The result of this calculation can be stated as follows,

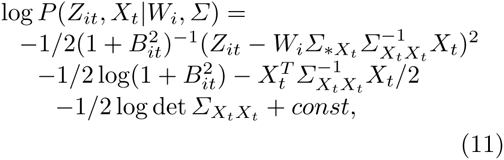
where the scalars 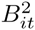 are defined by

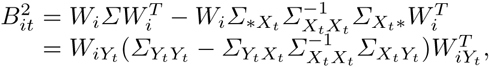
and the subscripted notation for *Σ* refers to the parts of *Σ* corresponding to the neuronal inputs identified in *X_t_* and *Y_t_*. For example, 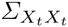 refers to the submatrix of *Σ* composed of all elements of *Σ* located at the intersection of the rows and the columns identified by *X_t_*. Similarly, 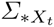 is the rectangular submatrix of *Σ* containing all the columns corresponding to the observed inputs *X_t_*, and 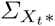 is a similar rectangular submatrix of all the *X_t_*-rows of *Σ*.

Eq. (11) allows us to obtain the final expression for the expected log-likelihood *l*(*W_i_, Σ*),

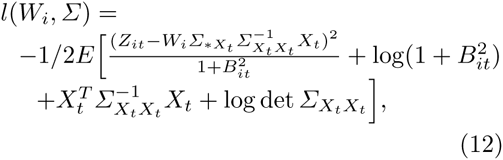
where the expectation is again with respect to the true distribution of the observed inputs and outputs *X_t_* and *Z_it_*. Consider now the expected log-likelihood 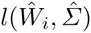 for an estimate of the parameters 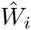 and 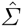. We first take the average in Eq. (12) over all *X_t_* such that the set of neurons contained in *X_t_* is fixed. This allows us to rewrite 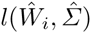 in the following form,

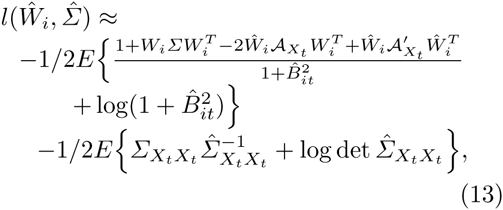
where the matrices 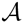 and 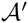 are defined by,

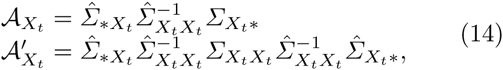
and *W_i_* and *Σ* are the true connection weights and the true covariance matrix, respectively, and where the remaining average is over the different subsets of observed neurons *X_t_*.

It can be verified by a direct inspection that 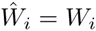 and 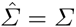 achieves the global maximum of Eq. (13) by separately maximizing the expressions under both expectation values for every *X_t_*, that is, 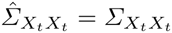 and 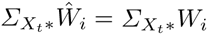. Moreover, this global maximum is unique whenever the covariance matrix *Σ* is nonsingular and the submatrices 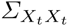 and 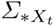 for the sampled *X_t_* cumulatively (but separately) tile the entirety of the matrix *Σ*. This is because the collection of individual-*X_t_* optimality conditions 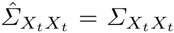 and 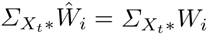 in this case uniquely specifies *Σ* and 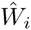.

However, we also find that Eq. (13) may contain local optima different from the global maximum. For example, a second local “mirror” optimum can be found at 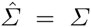 and 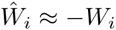, when the fraction of observed neurons in *X_t_* is small, Figure 1. This mirror optimum can be shown to arise from the cancellation of the variations of the terms 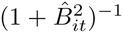 and 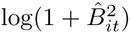 in Eq. (13). The difference in the expected log-likelihoods of the true and the mirror solution tends to zero with the the average fraction of the observed neurons *X_t_*, *p*, as *Δl* = *O*(*p*^2^).

**Fig. 1.**
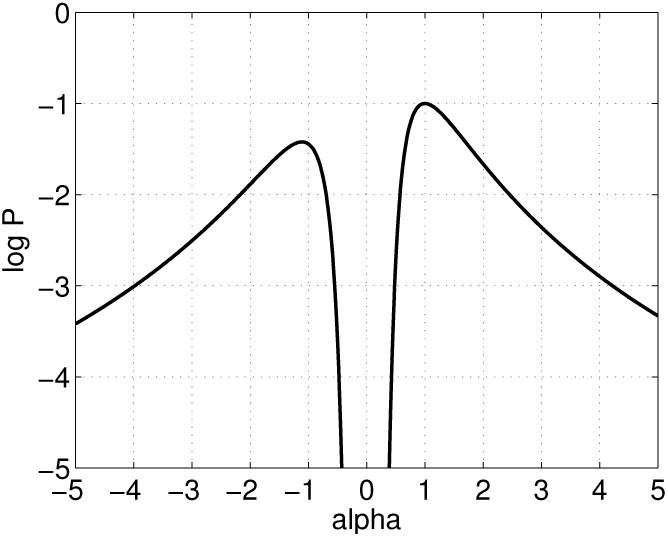
An example of 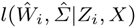 for a choice of *W_i_* and *Σ* = *I* along the solution ray 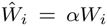, 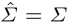. The plot shows that *α* = 1 is not a unique local maximum. The second “mirror” optimum is located at *α ≈* −1.

The conditions above give the sufficient conditions for the consistency of the MLE in the considered neuronal activity model. We point out that the condition of tiling of *Σ* by 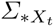 and 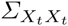 can be restated as the condition that the set of all observed neuronal subsets *X_t_* cumulatively covers the entire range of possible inputs *j* = 1 … *N* (that is, 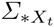 tiles *Σ*), and the set of all *X_t_* × *X_t_* covers the range of all possible input-input pairs {(*j, jʹ*) : *j, jʹ* = 1 … *N*} (that is, 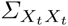 also tiles *Σ*). Furthermore, we can note that the fact of the full coverage of *Σ* by the collections of 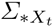 and 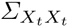 is important and not the manner in which it is achieved. Thus, any observations organization that can provide such a coverage will be equally capable of uniquely constraining 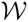. For example, a plausible strategy for estimating the complete connectivity matrix 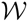 in these settings can be to image the activity of all individual neuronal pairs 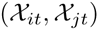 and 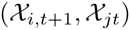, one pair at a time, in any order.

#### 2.2.3 The hidden inputs bias in the linear neural activity model

The expected log-likelihood *l*(*W_i_, Σ*) calculated in Section 2.2.2 can be used to provide an explicit example of the hidden inputs problem. Specifically, we consider the situation when the set of the observed input neurons *X_t_* is kept constant throughout the experiment, *X_t_* ≡ *X*. In this case, we can remove in Eq. (13) the expectation value with respect to the different sets of neurons *X_t_*. Performing the variation with respect to 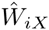 then yields the following equation for 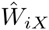,

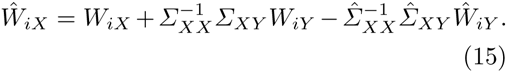

Eq. (15) shows that, if one has access to the incoming connection weights of the hidden neuronal population, 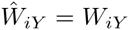, and the correct covariance matrix, 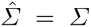, then it is possible to estimate the correct connectivity weights *W_iX_* even without observing the hidden neuronal activity. However, if hidden contributions are ignored, for example, by setting 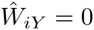 or 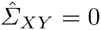, then the hidden inputs induce a bias in the estimated neuronal connectivity, which we obtain here explicitly as,

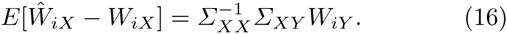

#### 2.2.4 The variance of the shotgun connectivity estimator in the linear neuronal activity model

We can evaluate the variance of the shotgun ML connectivity estimator in the linear neuronal activity model by calculating the Laplace approximation in Eq. (13) around 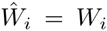. In this case, denoting 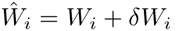, we find with respect to 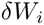,

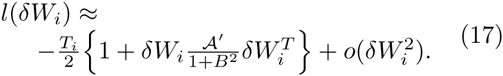

From Eq. (17), we read out the variance of the ML estimator 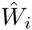 as,

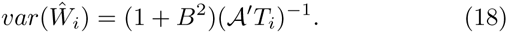

For weakly correlated case, *Σ* ≈ *I*, this can be reduced to a much more intuitive expression,

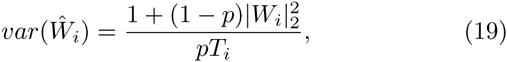
where *p* is the fraction of the neurons contained on average in one observation *X_t_*, and |*W_i_*|_2_ is the 2-norm of the input connectivity row-vector *W_i_*.

The above result indicates that the posterior variance of the shotgun connectivity estimator grows with the square of neuronal connectivity as well as the size of the unobserved neuronal populations. In addition, the estimator variance grows as 1/*pT_i_* as *p* drops.

As we will see in Section 3, this result has a particularly simple interpretation: The statistical error in 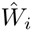 is introduced by the intrinsic noise *ϵ* as well as the uncontrolled variability of the hidden inputs 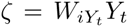. The reduction in that error for a given weight *w_ij_*·occurs in the proportion to the number of the observations in which the output of neuron *i* and the input of neuron *j* are simultaneously observed, *pT_i_*. If both the output neuron and the input neuron are observed randomly with probability *p*, as in the original shotgun proposal, then the estimator error respectively is,

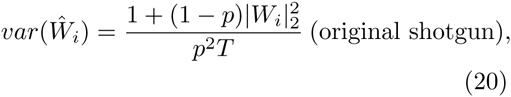
where *T* is now the total number of the observations and *p*^2^*T* is the number of the observations of different input-output neuronal pairs.

#### 2.2.5 The general correctness of the shotgun ML connectivity estimation in the linear neuronal activity model

Theorem 1 proved in Section 3.1 can be used to put in context the analytical results of Section 2.2.2. In particular, Theorem 1 shows that, if a neuronal population activity model parameter 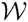 can be brought into a unique correspondence with a set of partial neuronal activity distributions 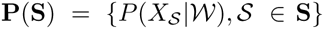, where *S* stands for different subsets of observed neurons and 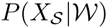 is the distribution of their activity, then such parameter can be uniquely estimated given any set of partial neuronal activity observations containing **P**(**S**). We can rephrase this statement to say that, if the set **S** is such that different model parameters 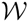 always identify distinct subpopulation activity distributions in **P**(**S**), then MLE provably converges to the true 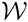 on the set of observations {*X_s_*, *S* G **S**}.

We can use this theorem to put in context the result of Section 2.2.2. In particular, by multiplying Eq. (2) on the left with 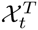 and averaging over time, we can obtain

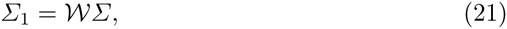
where 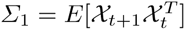 and 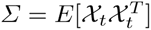. As long as the matrix *Σ* is nonsingular, Eq. (21) establishes a unique correspondence between the neuronal connectivity matrix 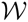 and the correlation matrices *Σ*_1_ and *Σ*. That is, if two connectivity matrices are not equal, by virtue of Eq. (21) they require two different pairs of the correlation matrices *Σ*_1_ and *Σ*. Since a given probability distribution implies a definite value of the correlation matrix, this establishes the desired correspondence between 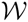 and the set of neuronal activity distributions 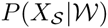.

More specifically, we conclude that in the linear neuronal activity model different connectivity matrices 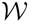 necessarily require different sets of neuronal activity probability distributions 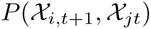 (defining the matrix *Σ*_1_) and 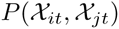 (defining the matrix *Σ*). By Theorem 1, any set of partial activity observations that fully specify these two sets is sufficient to uniquely identify the full 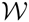. This statement is identical to the result of Section 2.2.2: providing all distributions 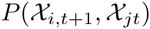 and 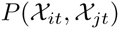 is equivalent to fully tiling *Σ* with the submatrices 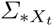 and 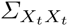.

Finally, we recall the note of Section 2.2.2 that, although the input-output model defined by Eq. (5) needs to be specified by two matrix parameters 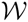 and *Σ*, the parameter *Σ* in fact is not independent, if considered within the scope of Eq. (2). For example, for the stationary distribution of Eq. (2) 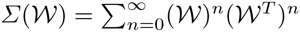 It can be shown then in principle that the single set of the distributions 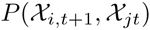 suffices to completely constrain the full model 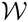.

Thus we note that, either non-deterministic sampling protocols or otherwise reasonable deterministic protocols that change which subpopulation is imaged more slowly than the recording device’s temporal resolution, can allow estimating all distributions 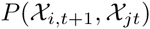 and 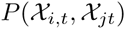, thus satisfying the sufficient conditions obtained in this section and in Section 2.2.2.

#### 2.2.6 The sufficient conditions for the correctness of the shotgun ML connectivity estimation in exponential generalized linear models of spiking neural activity

Of special interest to applications in basic neuroscience is the so called spiking generalized linear model (GLM) of neuronal activity with exponential nonlinearity (Brillinger, 1988; Rigat et al., 2006; Pillow et al., 2008). In greater details this model is described in this paper in Section 2.4.2. For this model, we obtain explicitly important results regarding the correctness of the shotgun ML connectivity estimation in the practically interesting case of the large number of neurons *N*.

More specifically, we examine the spiking exponential-GLM described by a nonuniform Poisson spiking process with the instantaneous spiking rate

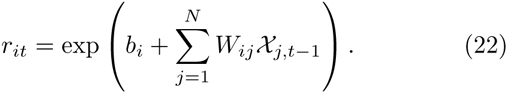

Note that here the probability of neuronal spiking depends only on the previous neuronal population’s state 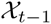. This situation is simpler to analyze, while it remains sufficiently general so that the GLM in the form given by Eq. (40-41) can be reduced to Eq. (22) by concatenating into 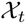 the neural spikes from many past times. Each 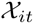 is binary, either 1 or 0, corresponding respectively to spike or no-spike of neuron *i* at time t. The standard log-likelihood is,

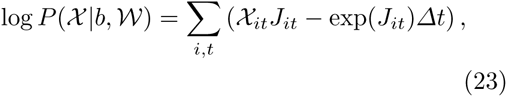
where

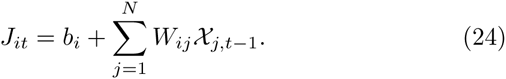

In the case when *N* is large, the law of large numbers states that the sum in Eq. (24) can be approximated under rather general conditions by a Normally distributed random variable with the mean 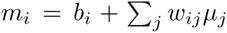 and variance 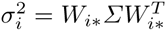, where 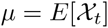 and 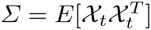 is the same-time covariance matrix, as before. The sum over the second term in Eq. (23) then can be replaced with

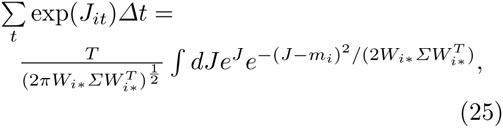
which clearly only depends on *m_i_* and *Σ*, aside from the sought model parameters *b* and 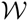. The sum over the first term in Eq. (23) is explicitly

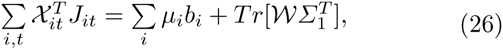
where 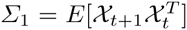 is the time-shifted covariance matrix, as before. We can conclude, then, that the log-likelihood of the exponential-GLM in the limit of large number of neurons is a function of only *μ*, *Σ*, and *Σ*_1_,

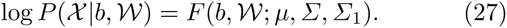

By Theorem 1 in Section 3.1, then, any set of partial observations completely constraining *μ*, *Σ* and *Σ*_1_ is sufficient to uniquely identify the full 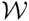. That is, any set of observations specifying all distributions 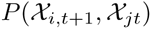 and 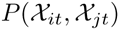 suffices to uniquely constrain 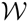, insofar as any two exponential GLM are distinguishable on the full set of observations.

Note that this also provides an extremely handy method for estimating 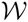 (Soudry et al., 2015). In particular, once *μ*, *Σ* and *Σ*_1_ had been calculated from any observations, 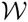 can be found by maximizing

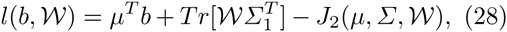
where

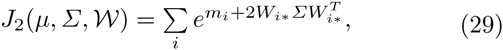
and the column-vector 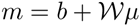.

We can also note, as before, that the parameters *μ*, *Σ* and *Σ*_1_ are not all independent and, in particular, knowing *μ* and *Σ*_1_ one can expect that *Σ* can be extracted from the dynamics equations of the model. Then, 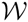 can be found by using

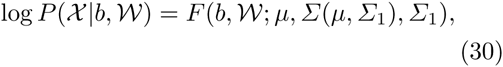
whereas the measurement of all 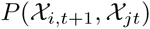 already suffice to determine 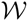. However, in that case one needs to explicitly obtain the solution for *Σ*(*μ*, *Σ*_1_), which may be more challenging in practice than simply measuring *Σ* separately.

#### 2.2.7 The sufficient conditions for the correctness of the shotgun ML connectivity estimation in general generalized linear models of spiking neural activity

Here, we extend the result of the previous section to general spiking generalized linear neural activity models, where the spiking rate is defined by

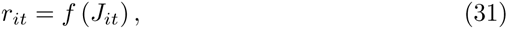
where *J_it_* is given by Eq. (24). The standard observations’ log-likelihood now is given by,

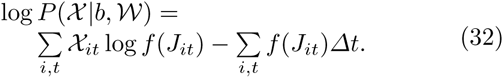

As before, in the limit of large *N* the distribution of *J_it_* can be approximated using Gaussian. Then, we can rewrite the log-likelihood (32) as

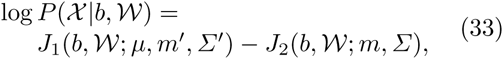
where

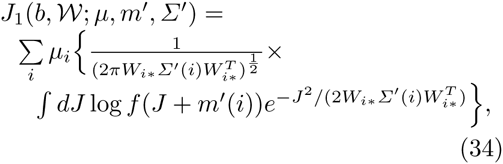
and

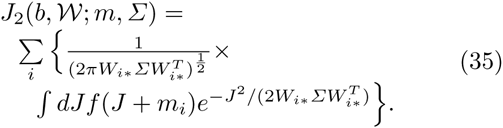

Where *m*, *μ* and *Σ* were defined in Section 2.2.6 and *mʹ*(*i*) and *Σʹ*(*i*) are,

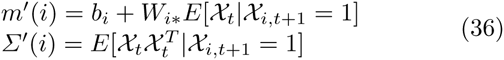

By Theorem 1 in Section 3.1, these imply that the set of distributions 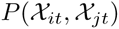, 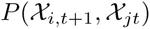 and 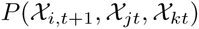 is sufficient to identify 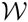.

### 2.3 The numerical solution of the shotgun connectivity estimation problem in general causal models of neuronal population activity

#### 2.3.1 The general causal model of neuronal population activity

The problem of estimating the effective connectivity matrix 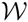 in realistic settings will require numerical solution. In this section, we develop a numerical approach for solving this problem using the Expectation Maximization algorithm. In the following, we assume throughout that the activity 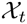 of the neuronal population in question can be modeled by a general Markov model defined by the relationship

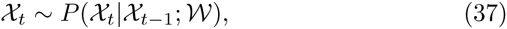
where 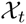 is the activity of the neuronal population at time *t*, defined to be conditional on the state of the neuronal population at time *t* − 1, 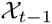, and some parameter 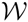. The linear neuronal activity model, Section 2.2, is evidently in this form. However, Eq. (37) is more general and covers a great variety of other important statistical approaches for neuronal activity modeling, with a particularly important case being that of the generalized linear model of neuronal activity, discussed in greater detail in Section 2.4.2.

#### 2.3.2 Expectation Maximization algorithm for the shotgun connectivity estimation problem in general neuronal activity models

We formulate the problem of connectivity estimation from the shotgun sampling data as the estimation of 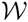 in model (37) over a set of partial observations of the neuronal activity 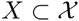. The standard method for solving such a parameter estimation problem in the presence of missing data is the Expectation Maximization (EM) algorithm (Dempster et al., 1977). To recap briefly, the EM algorithm produces a sequence of parameter estimates 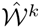 with uniformly increasing likelihoods 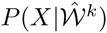, guaranteeing at least a locally-maximum likelihood estimate of 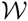. The sequence 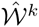 is produced by iteratively maximizing the functions

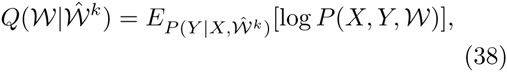
where 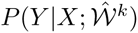 is the posterior distribution of the hidden neuronal activities given the available observations *X* and the current estimate 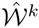. This maximization is typically realized by constructing *M* samples of the missing data *Y^l^* from 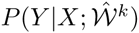 and then calculating 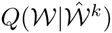 as 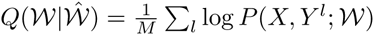.

To apply the EM algorithm here, we reformulate Eq. (37) as a Hidden Markov Model (HMM) in which the neuronal population activities, *Y_t_*, are treated as hidden states and the activities *X_t_* are treated as observations. We first construct the sample of the unobserved neuronal activities *Y*. Given the above definitions, this problem can be stated as the sampling of the hidden state sequences *Y_t_* from a HMM constrained on the observations *X*, and solved efficiently using the standard forwardbackward algorithm (Rabiner, 1989; Paninski et al., 2010).

Assuming the sample 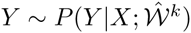 had been constructed, we evaluate 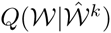 by using,

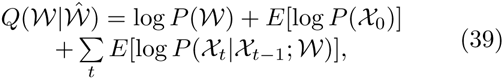
where the expectation values are with respect to the produced sample of *Y*. This function can be calculated straightforwardly and also can be made convex with a suitable choice of the causation in Eq. (37)—for example, by using a log-concave rate function *f* in Eqs. (40-41) (Paninski, 2004; Paninski et al., 2010). If the latter is achieved, the maximum of 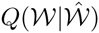 can be found efficiently with the help of standard gradient descent algorithms.

A detailed description of the implementation of the EM algorithm for model (37) is given in Appendix A.

### 2.4 Numerical simulations

#### 2.4.1 Numerical simulations of the shotgun connectivity estimation in the linear neuronal activity model

As a check of the calculations in Section 2.2.2, we performed numerical simulations of the shotgun connectivity estimation in the linear neuronal activity model. The model was simulated using the definition given by Eq. (5) for a single output neuron and a population of neuronal inputs. The input connection weights *W* were chosen uniformly at random on the interval [0, *W_max_*], with a connection probability *s*. The activities of the input neurons *X_t_* and *Y_t_* were drawn from a multivariate normal distribution with zero mean and a covariance matrix *Σ*, randomly generated from uniform distribution on [0, 1] and normalized to unit variances. More specifically, the elements of *Σ* were initially independently chosen from uniform random distribution on the interval [0, 1]. Then, thus obtained matrix was symmetrized by using *Σ* := *Σ* + *Σ^T^* and offset by using 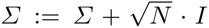, to guarantee the positive definitiveness (Furedi and Komlos, 1981). Finally, each raw and each column of *Σ* were divided by the square of the respective diagonal element, 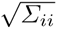, to normalize *Σ* to unit variances *Σ_ii_* = 1. The observations were simulated by continuously observing the activity of the output neuron and sampling the activities of the input neurons using the block-wise round-robin strategy, discussed in Section 3.2. The original connectivity was numerically recovered from thus simulated data using an implementation of the EM algorithm in Matlab, following the discussion of Section 2.3.2, with the sample size of the hidden neuronal activities set at *M* = 100.

#### 2.4.2 Numerical simulations of the shotgun connectivity estimation in the generalized linear neuronal activity model

In order to test the shotgun connectivity estimation in more realistic settings, we performed the numerical simulations of the shotgun connectivity matrix estimation using several model cortical neuronal networks, described using the generalized linear model (GLM) (Brillinger, 1988; Rigat et al., 2006; Pillow et al., 2008).

GLM is a particularly general and powerful class of statistical models of neuronal population activity, described as a nonuniform Poisson neuronal spiking process,

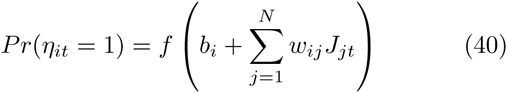

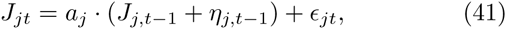

Here, *η_it_* is a binary variable specifying whether *i*-th neuron fired at time *t*, *f* is a nonlinear firing rate function, *b_i_* is a scalar offset, and *w_ij_*·are coupling weights. The driving currents *J_jt_* are defined as autoregressive processes with decay constants *a_j_* and a normal noise 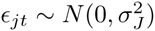. More than one autoregressive current may be defined per neuron, allowing the modeling of complex neuronal responses as the combinations of *J_jt_*’s with different decay constants. The diagonal weights *w_ii_* model the self-dependencies in neuronal activity, such as refractory period, bursting, etc., whereas the off-diagonal weights *w_ij_* model the interactions between neurons. Generalized linear models had been successfully applied in the literature to model the statistical properties of individual neurons as well as that of large neuronal populations (Brillinger, 1988; Chornoboy et al., 1988; Brillinger, 1992; Plesser and Gerstner, 2000; Paninski et al., 2004; Paninski, 2004; Rigat et al., 2006; Truccolo et al., 2005; Nykamp, 2007; Kulkarni and Paninski, 2007; Pillow et al., 2008; Vidne et al., 2009; Stevenson et al., 2009). The model given by Eqs. (40-41) falls under the general definition given by Eq. (37) after the identification 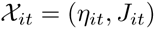.

In this paper, we simulated two GLM model neuronal networks—a small toy model of a synfire neuronal circuit and a larger realistic weakly coupled network of cortical neurons.

The synfire network was simulated in order to demonstrate the resolution of the canonical common inputs situation in the shotgun approach. The synfire model was created as an all-excitatory network of *N* = 10 neurons with strong feed-forward connectivity, as described in Figure 4. All connection strengths were chosen as a constant *W_syn_*, selected so that the probability of a post-synaptic neuron spiking conditional on a spike of a connected pre-synaptic neuron was approximately 80%. The model was then simulated using Eqs. (40-41) and the parameter definitions in Table 2. The neuronal activity was collected using the block-wise round-robin sampling, described in Section 3.2.

**Table 1.** The parameters of the linear neuronal activity model used in the numerical simulations in this paper. 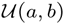 is a uniform probability distribution on [*a, b*].

|  |  |
| --- | --- |
| Number of input neurons, $N$ | 100 |
| Input weights distribution, $W$ | $\mathcal{U}(0, W_{max})$,<br>$W_{max} = 2$ |
| Connection probability, $s$ | 10% |
| Observations sparseness, $p$ | 20% |
| Sample size, $T$ | 500 samples |

**Table 2.** The parameters of the toy GLM synfire model used in the numerical simulations in this paper.

|  |  |
| --- | --- |
| Number of neurons | 10 |
| Base log-firing rate ($\exp(b_i)$) | 15 Hz |
| Average firing rate | 33 Hz |
| Simulation time step | 1 msec |
| Max observation duration | 50 sec |
| Neurons per imaging block | 20% |
| Excitatory conn. strength ($w_{ij}$) | 13 |
| Excitatory PSP rise time | 1 msec |
| Excitatory PSP decay time | 1 msec |
| Refractory time | 1 msec |

The model of a weakly coupled cortical neuronal network was adopted here from (Mishchenko et al., 2011), and was intended to demonstrate the shotgun connectivity estimation in a more realistic setting. The model closely reproduced the experimental data about local cortical neuronal circuits available in the literature (Braitenberg and Schuz, 1998; Gomez-Urquijo et al., 2000; Lefort et al., 2009; Sayer et al., 1990). More specifically, the neuronal population was created as 80% excitatory and 20% inhibitory neurons. The neurons were connected with each other randomly and homogeneously with the probability of 10%. The Dale’s law was respected. The strength of the excitatory connections was set using the peak excitatory post-synaptic potential (PSP) values randomly chosen from an exponential distribution with mean 0.5 mV (Lefort et al., 2009; Sayer et al., 1990). The strength of the inhibitory connections was set using a similar exponential distribution with the mean chosen to balance the average excitatory and inhibitory inputs in the network (Abeles, 1991). All neurons had the refractory periods of 1 msec, enforced in Eqs. (40–41) via the self-currents *J_it_* with the decay time-constants 1 msec and *w_ii_* = −100. Individual PSPs were modeled using the alpha function (Koch, 1999), described as the difference of two exponentials with a rise time of 1 msec and a decay time of 10 to 20 msec (Sayer et al., 1990). Given the simulations time step of 1 msec here, such PSPs can be described more precisely as an instantaneous jump followed by an exponential decay of 10 to 20 msec, as described by Eq. (41). The spiking activity was simulated at 1 msec time step ignoring the interneuronal conduction delays, negligible for spatially compact neuronal circuits. The neuronal activities were downsampled at 100 Hz, in order to simulate the observations using a slow imaging technique such as calcium imaging. Generated neuronal activities were collected using the block-wise round-robin sampling strategy, Section 3.2. The detailed parameters used in the simulations are listed in Table 3.

The connectivity matrix was estimated from the simulated data using a Matlab implementation of the EM algorithm described in Appendix A. The sample size of the hidden neuronal population activity was taken as *M* = 50 for the cortical neuronal networks and *M* =150 for the synfire network. Only the connectivity matrix weights were estimated, taking the PSP time-constants as known. This was done to separate different sources of errors in the estimation, understanding that the focus here is on the problem of the shotgun connectivity inference. The PSP time-constants, more generally, can be set uniformly in the population from physiological data without a significant impact on the connectivity estimation, or also included into the EM procedure. For more in depth discussion of this issue see (Mishchenko et al., 2011).

The EM algorithm was executed on a 4 dual-core i7 processor desktop computer with 8GB of RAM memory. The numerical estimation problems could be generally solved in a reasonable amount of time, however, we found that the necessity to keep up to *M* samples of the complete activity of the hidden neuronal populations imposed drastic requirements on RAM memory. In particular, for the GLM cortical neuronal networks described here and using the said workstation, we could solve at most the models with *T* = 3000 seconds of neuronal activity data and *M* = 50 examples of the hidden activity data. The above configuration, thus, set the limits on the numerical experiments performed in this paper.

More generally, in order to set the parameter *M* of the number of samples used to model the posterior distribution of hidden neuronal activities, we tried different *M* such as *M* = 10, 50, 100 and 150. The value of *M* = 50 was the highest value that we could try on the above mentioned computer configuration for the GLM neuronal networks. The value *M* = 150 was the highest that we could try for the synfire model. Generally, we found that *M* = 10 sufficed to obtain a solution, however, the noise in that solutions was high and correlated but disconnected neurons frequently were identified as connected (in the case of the synfire model). Using a larger hidden activity sample *M*, thus, appeared to be necessary. *M* = 50 appeared to be sufficient to achieve satisfactory reconstructions for the weakly correlated cortical neuronal models, but appeared to be lacking in the case of the strongly correlated synfire model. *M* above 100 appeared to be satisfactory in the case of the weakly correlated cortical neuronal network as well as the synfire model.

## 3 Results

### 3.1 The correctness of the shotgun neuronal connectivity estimation

One of the biggest challenges of the functional connectivity estimation in neuroscience remains the presence of unobserved or hidden inputs in neuronal population activity data. The shotgun connectivity estimation is a promising approach for alleviating this problem, consisting in imaging a large neuronal population using small groups of random neurons and reconstructing the complete connectivity matrix from such partial observations.

In Materials and Methods, we study analytically the problem of the shotgun connectivity estimation in a linear model of neuronal population activity. We show that the shotgun estimation can be always posed as the problem of estimating the input connectivity of one “output” neuron at a time, given a partially observed population of neuronal inputs. This is because in typical network models of neuronal activity the activity of neurons is conditionally independent given the activity of their presynaptic neuronal populations and the respective connection weights.

In the considered linear neuronal activity model, we explicitly calculate the observations likelihood 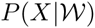 and demonstrate that the MLE in this case converges to the correct complete connectivity matrix as long as all possible input-input and input-output pairs of neurons are observed together in at least a fraction of the observations. We also explicitly derive the hidden inputs bias and the variance of the shotgun estimator in this model.

It is possible to further extend these results to more general settings. For that, we assume first that the activity of a neuronal population, 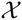, can be described by a general parametric model probability density 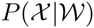, given a parameter 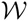. We say that an estimator 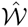 is consistent as long as 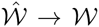 as the sample size tends to infinity. Given a set of partial neuronal activity observations 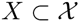, we recall from the asymptotic estimation theory that the MLE is guaranteed to provide a consistent estimator as long as the observations distribution 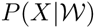 possesses no observation-indistinguishable parameter sets. That is, 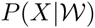 is such that no two distinct parameters 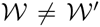 result in identical distributions 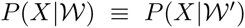. The above condition is commonly referred to in statistics as the identifiability property.

**Table 3.**
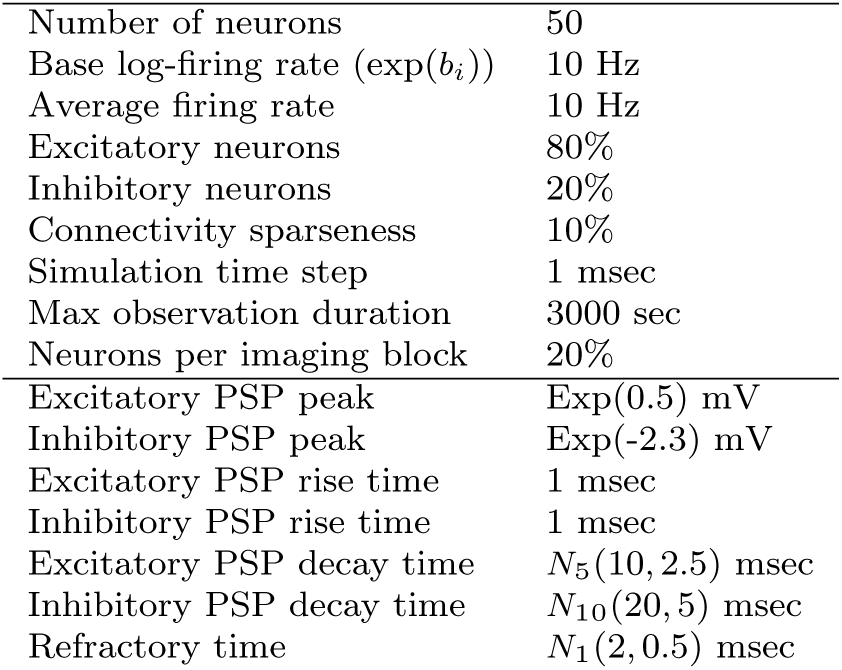
The parameters of the realistic GLM cortical neuronal network used in the numerical simulations in this paper. Exp(λ) is the exponential probability distribution with mean λ, and *N_p_*(*μ, σ*) is the truncated-normal probability distribution with mean *μ*, standard deviation *σ*, and lower bound *p*.

The above observation can be made most readily by considering the average observed-data log-likelihood maximized by the MLE. More specifically, let us consider a collection of neuronal population activity observations {*X*^(^*^k^*^)^, *k* = 1 … *n*}. This can be understood as a collection of repeated experiments or a collection formed from different segments of the same experiment, corresponding to its different time-intervals.

In the MLE, we maximize the log-likelihood of the observed data, whereas by the observed data here we understand the collection of all neuronal activity observations *X*^(^*^k^*^)^. The average observed-data log-likelihood in the limit of large n converges in probability to

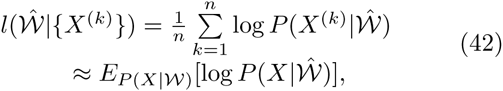
where the last line identifies the expected log-likelihood function 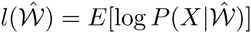, and the expectation is with respect to the true distribution of *X*, 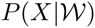. Well known Gibbs inequality then tells us that Eq. (42) achieves its global maximum when and only when 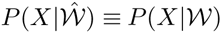 for all *X*. The above standard argument leads to nontrivial insights about the shotgun neuronal connectivity estimation below.

From the perspective of the identifiability of the connectivity estimation problems in neuronal activity models with the lack of complete observations, we first point out that the problem of hidden inputs in that setting arises because the identifiability condition becomes broken. That is, different models of hidden neuronal connectivity can produce the same distribution of the observed data. In the classical example of the hidden inputs problem, for example, a direct connection is inferred between two unconnected observed neurons because a correlated input fed into these neurons by a third neuron also can reproduce the data collected on the observed neurons. The violation of identifiability leads to multiple network models being able to reproduce the same empirical distribution 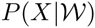 and, therefore, achieve the global maximum of the log-likelihood (42). However, the key observation at this stage is that the true connectivity 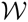 still remains a ML solution. Thus, it is not that the hidden inputs problem somehow skews the neuronal connectivity estimation. It is just that such estimation is no longer unique.

To examine this state of things in the shotgun estimation, we recognize initially that the expected log-likelihood (42) in this case should be re-written as,

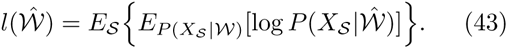

Here, 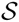 denotes a particular subpopulation of observed neurons and the external average is over such subpopulations 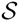 imaged during a shotgun sampling experiment. *X_S_* refers to the part of the neuronal population’s activity observed in 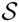, and 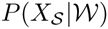 is the probability distribution of such activities.

Once again, we recall the observation above that the true connectivity matrix 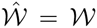 is necessarily a global maximizer of Eq. (43), by Gibbs inequality. More significantly, 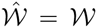 maximizes Eq. (43) by simultaneously maximizing all individual terms 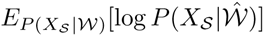, since evidently 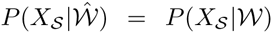 for all 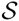 whenever 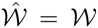. We now ask whether there can exist another model 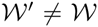 such that achieves the same global maximum. We note that, since 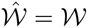 achieves the global maximum of Eq. (43) by simultaneously globally maximizing all 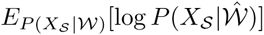, any other such maximizer of Eq. (43) must also have this property. In turn, this implies that any such 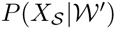 must match the marginal distributions 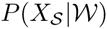 on all inspected 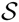.

We thus arrive at the following conclusion:

#### Definition 1

Assume that a statistical model of neuronal population activity 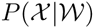 and a set of neuronal subpopulations 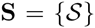 are such that for any two different model parameters 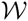 and 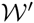 there exist at least one 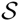 in **S** such that the distributions 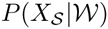 and 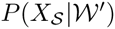 are not identically equal, where 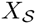 is the restriction of 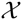 to 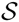. Then, we say that the model 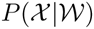 is uniquely identified by the set of distributions 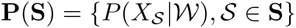.

#### Definition 2

Assume that for any neuronal subpopulation 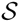 in a set of neuronal subpopulations **S** there exist a subpopulation 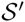 in a different set 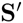 such that 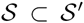. Then we say that 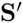 completely covers **S**.

#### Theorem 1

*Assume a statistical model of neuronal population activity* 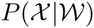 *is uniquely identified by a set of distributions* 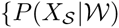, 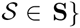*. Then, for any set of partial observations of the neuronal population activity in this model on a set of neuronal subpopulations* 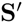 *that completely covers* **S**, *the ML estimator defined as the argmin of (43) is consistent*.

In other words, if the shotgun sampling **S**^ʹ^ covers all neuronal subpopulations 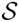 from the identifying set of partial neuronal activity distributions **P**(**S**), then the shotgun ML estimator is guaranteed to converge to the correct 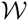. The condition on **S**^ʹ^ to completely cover **S** ensures that any 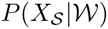 can be recovered from some 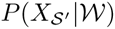 such that 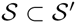, by marginalization.

Theorem 1 replaces the question of the consistency of the MLE in the shotgun sampling by the question of finding the identifying sets **P**(**S**) for a given neuronal activity model. It shows that, whenever an identifying set **P**(**S)** is covered by a sampling scheme, either using non-deterministic or deterministic protocol, the respective inference problem possesses the identifiability property, whereas in general such identifiability property for an inference problem with only partial observations data is unknown and, in fact, cannot be guaranteed even when the original full problem is identifiable. It is also worth noting that in Theorem 1 it is not important how exactly the mapping between the identifying sets **P**(**S**) and 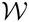 is constructed, or by using which statistics of the distributions in **P**(**S**) the parameter 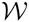 can be extracted and how. As long as such a mapping exists, the full parameter 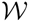 is recoverable by Theorem 1 by using the MLE.

It remains to establish the identifying sets of given neuronal activity models. In certain cases, such as in the linear neuronal activity model, the identifying set can be rather immediately established (Section 2.2.5), providing the sufficient conditions for the shotgun sampling to be successful. However, in general, different neuronal activity models require separate investigation into their identifying sets. From general arguments, we can conjecture that in a large class of network models of neuronal activity the set of all pair-wise input-output neuronal activity distributions such as 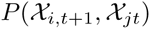 would suffice to form such a set, see below. However, in general the question of finding the identifying sets **P**(**S**) of a neuronal activity model is nontrivial, and even the existence of nontrivial identifying sets **P**(**S**) cannot be always guaranteed.

Specifically, consider an arbitrary probability density 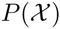 defined on a discrete grid of *d* points in *n* dimensions, with *d^n^* points in total. Such 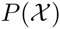 has *d^n^* − 1 free parameters. All marginal distributions of up to *n* − 1 dimensions together provide *n*(*d*−1) + *n*(*n* − 1)/2(*d*^2^ − 1) + *n*(*n*−1)(*n*−2)/3!(*d*^3^ − 1) +… < (1 + *d*)*^n^* − *d^n^* < *nd^n^*^−1^ linear conditions on 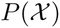. Therefore, even all marginal distribution 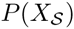 together are not sufficient to determine 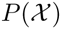 uniquely whenever *d* > *n*.

As a specific counter-example, let us consider a toy model 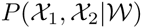 with 2 neurons in 4 possible states (*n* = 2 and *d* = 4). Specifying such a general model requires providing 4^2^ − 1 = 15 different probability values 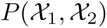. However, the two marginal distributions 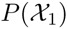 and 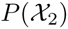 provide only 8 linear constraints on 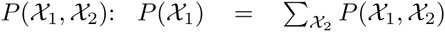 and 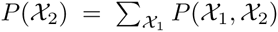. Clearly, an infinite number of 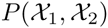 can be suggested perfectly matching these constraints for almost any 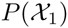 and 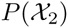. In this case, any partial observation of 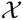 is insufficient—a complete observation is required.

We ran into the problems in the above example because the number of the degrees of freedom of the model probability distribution 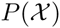 was simply too high. However, in network models of neuronal population activity 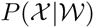 is generally a much more constrained distribution, which is typically completely specified by a single connectivity matrix 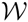 of *N*^2^ elements, where *N* is the number of neurons. Counting the degrees of freedom in this case implies that there can be at most *N*^2^ independent distributions in the set of all marginal distributions 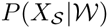. Choosing any subset of *N*^2^ independent such distributions, thus, will uniquely fix all other distributions and, therefore, the complete model 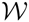.

It is natural to conjecture (but we cannot prove this now rigorously) that such a subset can be chosen in the form of the *N*^2^ inputoutput distributions 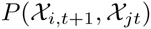. Other choices may also exist. For example, if the stationary distribution of the model (37), 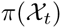, is such that a unique mapping is formed by 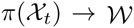, then a set of *N*^2^ independent same-time distributions such as 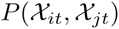 may do just as well for specifying 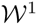^1^.

In summary, we suggest, based on the counting of degrees of freedom, that one can expect the set of the observations of all distinct inputoutput neuronal pairs 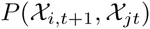 to be sufficient for recovering the complete neuronal connectivity matrix in general network models of neuronal population activity. At the same time, the above argument cannot constitute a rigorous proof. Therefore, we must present it here as a conjecture open for further investigation.

### 3.2 Alternative organizations of the shotgun neuronal population activity sampling

The results obtained in Section 3.1 have certain implications for the organization of the shotgun connectivity estimation experiments. We see that, to be able to reconstruct a complete neuronal connectivity, it is only important that the identifying set **P**(**S**) is covered by the observations, and the manner in which such coverage is provided is not important. Per the conjecture of Section 3.1, any imaging organization of a neuronal network that furnishes the observations of all input-output neuronal pairs in that network would suffices to uniquely identify the complete neuronal connectivity matrix.

A particularly advantageous and conceptually simple alternative organization of neuronal activity sampling from this perspective is the block-wise round-robin sampling illustrated in Figure 2. In this approach, a neuronal population is imaged as a series of contiguous blocks, one input and one output block imaged simultaneously for a set number of observations *Tb*. The blocks are moved through the population so that all possible combinations of the input and the output blocks are inspected.

The block-wise round-robin strategy can be straightforwardly implemented using the existing fluorescent microscopy tools by scanning two field-of-views of a confocal or two-photon microscope over the neuronal population. Another advantage of this strategy is a simpler numerical connectivity estimation problem.

**Fig. 2.**
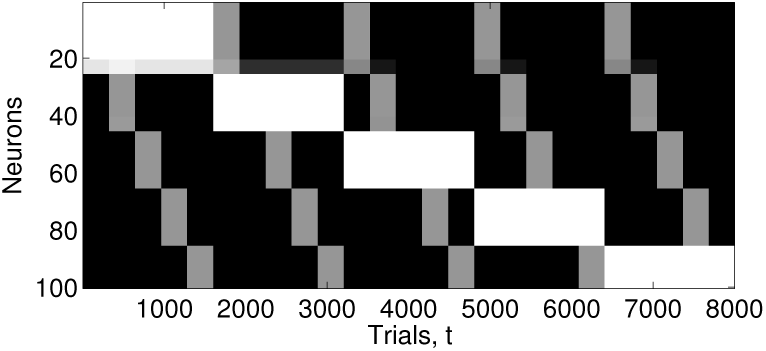
Block-wise round-robin neuronal population activity sampling strategy. In this strategy, the neuronal population is imaged as a sequence of contiguous blocks of input and output neurons. During one section of the experiment, the neurons in one input and one output block are observed simultaneously for a set of observations *T_b_*. All possible combinations of the input and the output blocks are observed over the entire experiment. The figure illustrates the block-wise round-robin sampling strategy applied to a hypothetical population of 100 neurons with 20 neurons observed per each input and output block. White color indicates the neurons in the output blocks and gray color indicates the neurons in the input blocks. In each observation, the activity of all marked neurons is observed simultaneously.

**Fig. 3.**
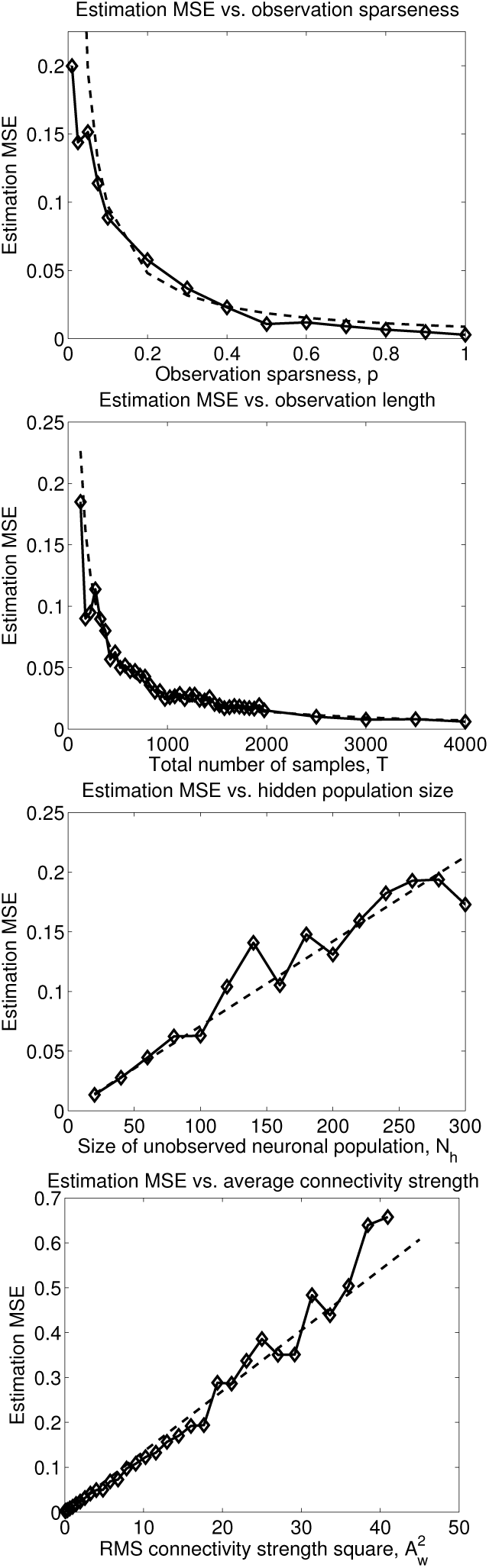
The properties of the shotgun connectivity estimator in relation to the missing data. From top to bottom, the posterior error of the shotgun estimator is shown in relation to the observations sparseness, the total number of observations, the size of the hidden population, and the rms average connectivity strength. The results of numerical simulations are shown with solid lines and the theoretical predictions are shown with dashed lines. The model parameters are described in Table 1.

Block-wise round-robin sampling strategy is sufficient for collecting all input-output as well as same-time pairs of neurons and, in that sense, is sufficient for the recovery of the complete neuronal connectivity in the sense of the conjecture about the identifying sets of network models of neuronal activity in Section 3.1.

### 3.3 The impact of the missing data on the shotgun connectivity estimation

In Section 2.2.4 we obtained some key properties of the shotgun connectivity estimator in the linear neuronal activity model. It is interesting to return to these results now, from the positions of the discussion of the last sections.

In particular, let us consider a segment of a block-wise round-robin neuronal activity sampling experiment with a fixed input and output neuronal blocks. We can roughly relate the activity of the observed neurons in this scenario as,

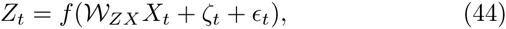
where *Z_t_* is the column-vector of the activities of the neurons in the output block, *X_t_* is the column-vector of the activities of the neurons in the input block, and we introduced a new random variable 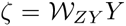, which represents the combined input of the hidden neurons into the neuronal outputs.

Posed from this perspective, the shotgun connectivity estimation appears now as the problem of estimating a block of the connectivity matrix 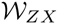 given the observations of the activities of all relevant input and output neurons, whereas the presence of the unobserved neurons enters solely in the form of an additional noise ζ. The noise ζ is both structured and correlated with *X*. The knowledge of that structure is required to successfully remove the associated bias from the estimates of 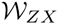. However, ζ also introduces additional statistical uncertainty into the estimates of 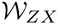. We can estimate the variance of ζ as 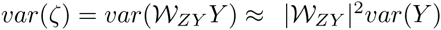, leading to a rough estimate of the variance of the estimator 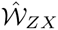 as,

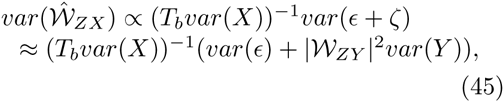
where 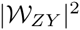 is understood as the average 2-norm of the rows of 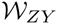. In the limit where the size of the hidden neuronal population is large, we can write that approximately,

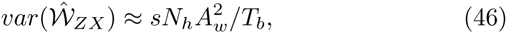
where *N_h_* is the number of neurons in the unobserved neuronal population, *A_w_* is the root mean square average of the (nonzero) neuronal connectivity weights, *s* is the sparsity of the connectivity matrix, and *T_b_* is the number of observations observing given input-output neuronal pairs.

Eq. (46) provides a useful approximation for the error of the shotgun connectivity estimator. We see that the primary source of that error is the uncontrolled fluctuations of the hidden inputs in the observed neurons. The error is affected by both the size of the hidden population and the strength of its coupling, and is reduced in the proportion to the total number of the observations of different input-output neuronal pairs.

### 3.4 Numerical simulations

In this section we demonstrate the recovery of the complete neuronal connectivity matrix from partial neuronal activity observations using simulated neuronal population activity data.

We first examine the problem of such a connectivity estimation in a small model synfire network of *N* =10 neurons, Figure 4. Synfire model represents one of the worst cases of the hidden inputs problem in the functional connectivity inference, as the correlations can propagate down the synfire chain over large distances and emulate false connections among distant neurons.

**Fig. 4.**
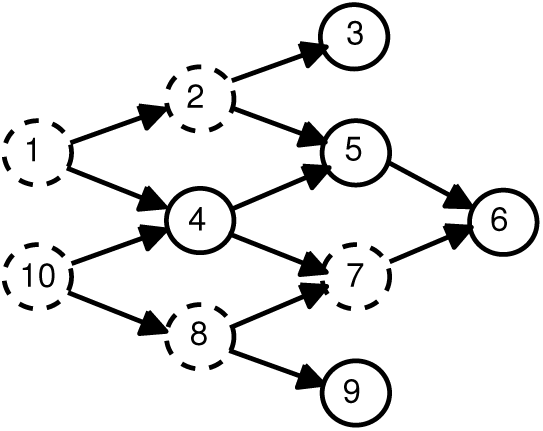
A model fragment of synfire neuronal circuit illustrating the resolution of the hidden inputs situation in the shotgun connectivity estimation.

We first illustrate the connectivity estimation in this model in the population of 5 neurons indicated in Figure 4 with solid circles. This corresponds to the typical neuroimaging situation, in which a fixed neuronal population is continuously observed whereas the rest of the neurons or their inputs are hidden. As a measure of connectivity, we present in Figure 5 the calculation of the time-shifted correlogram (of lag 1 bin) for the selected neuronal population, widely used in the literature as a functional connectivity measure, and a generalized linear model (GLM) connectivity matrix estimation. We observe that either the correlogram and the GLM connectivity matrix produce spurious connections in this neuronal population, namely, such seen between the neurons 4 and 9, 3 and 6, and 9 and 6.

**Fig. 5.**
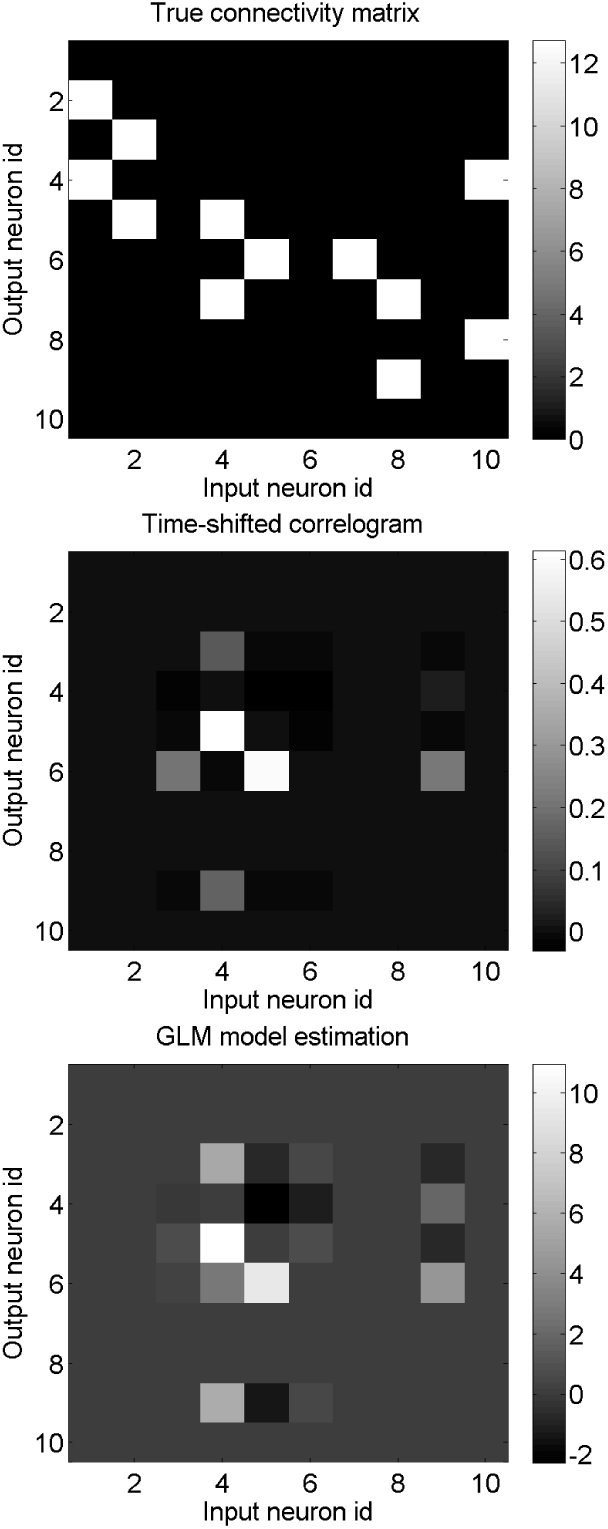
The hidden inputs problem in the model synfire circuit in Figure 4. The top panel shows the true connectivity matrix for the circuit in Figure 4. The middle panel shows the time-shifted correlogram calculated for the population if neurons indicated in Figure 4 with solid circles, from *T* = 10 seconds of observation. The bottom panel shows the GLM connectivity matrix estimation for the same population from *T* =10 seconds of neuronal activity data. The simulation parameters are as described in Table 2.

Next, in Figure 6 we perform the estimation of this neuronal connectivity matrix using the block-wise round-robin sampling strategy and the EM estimation algorithm described in Section 2.3.2. Despite the fact that at most 4 neurons are observed in this situation at a time, the complete neuronal connectivity matrix is recovered rather well. The strength of the false-positive connections between the neurons 4-9, 3-6 and 9-6 is also reduced by a factor of 2 to 3 compared to the reconstruction in Figure 5. At the same time, the obtained connectivity matrix is substantially more noisy and requires more observations to suppress that noise. Typically, we observe that the accuracy of the shotgun connectivity estimation with the observation time *T* is comparable to that using the full observations and the observations duration *p*^2^*T*, essentially in agreement with Eq. (46), the bottom panel in Figure 6.

**Fig. 6.**
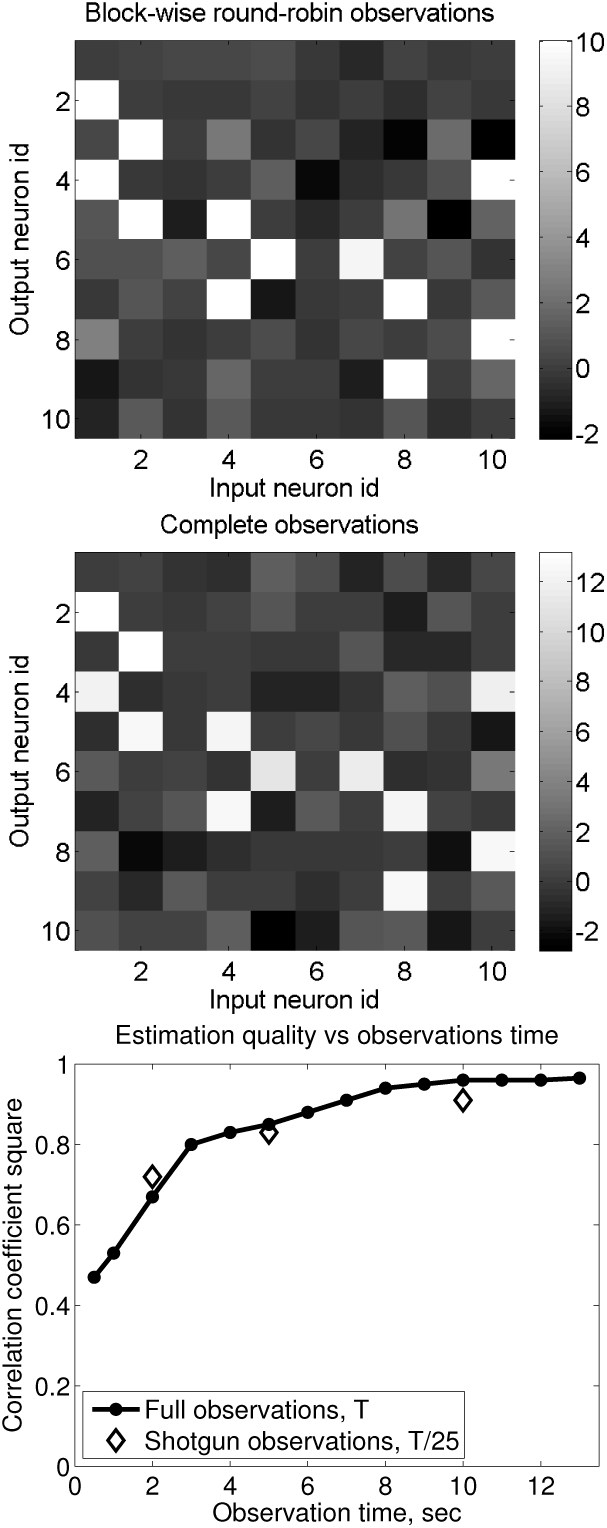
Estimation of the complete neuronal connectivity in the model synfire circuit in Figure 4 using the block-wise round-robin activity sampling. The top panel shows the result of the neuronal connectivity reconstruction using this approach and *T* = 50 seconds of neuronal activity data. The middle panel shows, for comparison, a similar reconstruction but using the complete observations and *T* =10 seconds of neuronal activity data. The bottom panel compares the quality of the estimated connectivity matrices for the block-wise round-robin approach (diamonds) and the complete data (solid line). The shotgun approach points are placed at the ’’equivalent” time calculated as *T′* = *p*^2^*T*. The simulation parameters are given in Table 2.

In Figure 7, we apply the shotgun approach to the estimation of the complete connectivity matrix in a model of realistic weakly coupled cortical neuronal network with *N* = 50 neurons. The shotgun estimation again allows recovering the complete neuronal connectivity matrix from partial neuronal population activity observations. Once again, the data size necessary to achieve a given accuracy is significantly greater than that required when using the complete neuronal activity. Specifically, if a good connectivity matrix reconstruction could be obtained in this simulation with all neurons observed from a total of *T* = 60 seconds of neuronal activity data, in *p* = 20% block-wise round-robin sampling case the imaging time necessary for a similar reconstruction accuracy approached T = 3000 seconds, the lower panel in Figure 7.

**Fig. 7.**
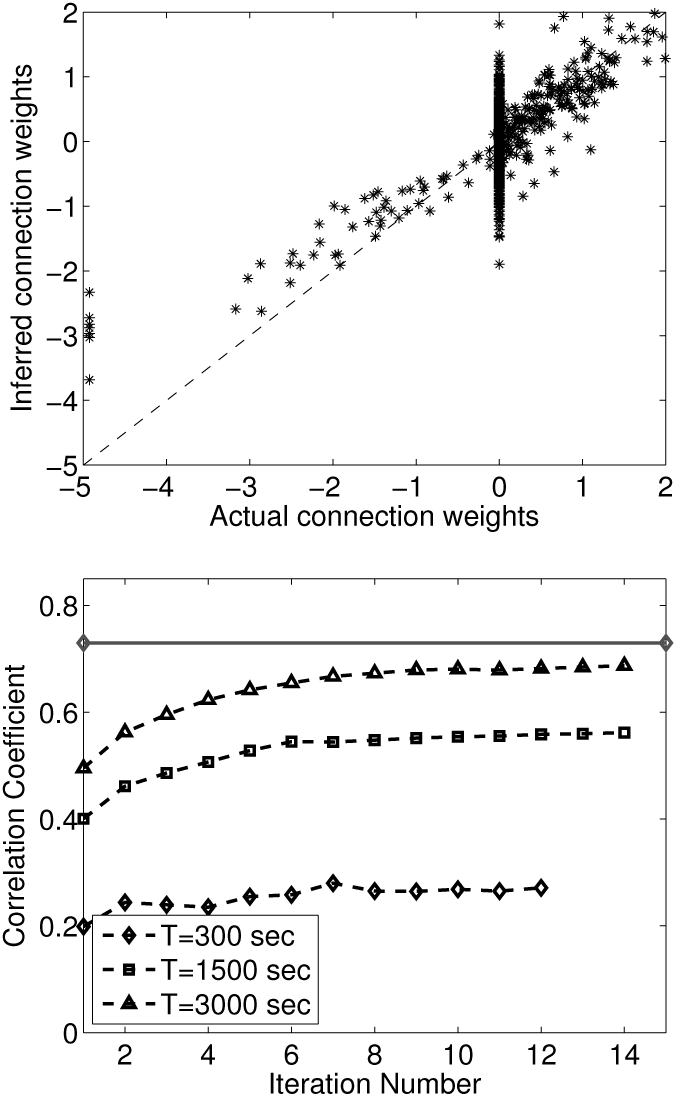
The shotgun connectivity estimation in a model of a realistic weakly coupled cortical neuronal network. The upper panel shows the reconstruction of the complete connectivity matrix using *p* = 0.2 block-wise round-robin neuronal activity observations in such network and *T* = 3000 seconds of neuronal activity data. The multiplicative bias seen in the reconstructed weights is the finite time-discretization bias discussed in (Mishchenko et al., 2011) (not corrected here). The reconstructed connection weights reproduce the true connectivity, however, are rather noisy. The bottom panel shows the correlation coefficient of the reconstructed connection weights vs the true connectivity as a function of the EM algorithm’s iteration number, for *T* = 300, 1500 and 3000 seconds of neuronal activity data. Solid gray line represents the baseline reconstruction produced using the complete observations and *T* = 60 seconds of neuronal activity data. The simulation parameters are given in Table 3.

## 4 Discussion and Conclusions

The shotgun sampling solution of the common inputs problem is a promising new approach for the functional estimation of the neuronal connectivity in large neuronal networks in the brain without requiring the simultaneous imaging of entire neuronal populations. By statistically estimating the neuronal connectivity from a collection of partial observations of different neuronal sub-populations, the shotgun sampling offers a possibility of recovering the connectivity matrix in realistically large neuronal circuits using limited imaging resources.

In this paper, we investigate analytically and in simulations the properties of such shotgun neuronal connectivity estimation. In Theorem 1 of Section 3.1 we establish the sufficient and necessary conditions for the complete connectivity matrix of a neuronal population to be recoverable from shotgun-type incomplete neuronal activity observations. In Section 2.2.2, 2.2.3 and 2.2.4, we discuss the shotgun connectivity estimation in a linear neuronal activity model more explicitly, and analytically derive its shotgun observations likelihood function, associated maximum-likelihood estimator, and the key properties of that estimator. In Section 2.2.5, 2.2.6 and 2.2.7, we explicitly establish the sufficient conditions for the correctness of the shotgun connectivity estimation in linear and exponential GLM neuronal activity models as well as in general spiking models of neuronal population activity, where the neuronal firing rates are described by a general nonlinear function of linearly summed inputs. The latter, in particular, covers a great variety of general network models of neuronal populations activity.

In the linear and the exponential-GLMs of neuronal activity in the limit of large number of neurons, we find that the shotgun estimation is guaranteed to produce the complete connectivity matrix whenever all possible input-output pairs of neurons as well as all the pairs of same-time neuronal activities had been observed in at least some parts of a shotgun sampling experiment. In these models, constraining all elements of the same-time and time-shifted co-variance matrices, 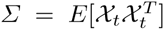 and 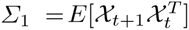 respectively, suffices to uniquely identify the full connectivity matrix as well as offers a simple way for deducing the connectivity matrix with modest computational effort, as discussed in Section 2.2.5 and 2.2.6. For general spiking GLM of neuronal population activity, we find that the set of observations of all neuronal activities in the form 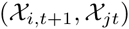 and 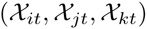 can be guaranteed to allow the recovery of the full connectivity matrix, as discussed in Section 2.2.7.

We find that, whenever the full connectivity matrix is uniquely identified by a set of observations, the observations likelihood function is as well guaranteed to have the correct connectivity matrix as a unique global maximum. At the same time, observations likelihood cannot be guaranteed to have a unique local optimum. Already in the case of the linear neuronal activity model, where the MLE solution in the full-observations case is globally and locally unique, we find the possibility of two local optima—the true solution 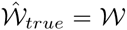 and the “mirror” solution 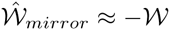, Section 2.2.2.

The mirror optimum-like solutions clearly will need to be avoided in practice. Simple heuristics may be able to achieve this objective in many cases, such as inspecting the likelihood of the “mirror” solution for any found locally optimal 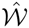. However, this strategy may not be plausible when the fraction of observed neurons *p* tends to zero, as we can show that the difference between the likelihoods of the true and the mirror solutions in this case quickly tends to zero. Alternatively, the resolution of the mirror-optima can be based on the use of a-priory information. Such a-priory information may include the identity of one or several excitatory or inhibitory neurons in the population or the relative abundances of the excitatory and inhibitory neuronal populations, which are rather easy to assess. By requiring given neurons in the reconstruction to be excitatory or by requiring a particular split between the abundances of the excitatory and inhibitory neuronal populations, the sign of the solution can be fixed trivially.

We find that the lack of the observations of the entire neuronal population in the shotgun estimation leads to an increase in the statistical error of the connectivity estimator, warranting a respective increase in the imaging time necessary to suppress that error. We find that the required shotgun sampling data size scales proportionally to the number of neurons in the unobserved neuronal populations and the average square neuronal connectivity strength. The data size also increases as the inverse square of the fraction of neurons in the shotgun observations. This scaling is inopportune for the reconstructions of neuronal connectivity where the fraction of observed neurons will remain small.

In Section 2.3.2, we propose an exact numerical approach for solving the shotgun connectivity estimation problem in general settings, and demonstrate its applications to the estimation of the complete neuronal connectivity matrix in different model neuronal circuits, including linear neuronal models, a spiking synfire network, and a small realistic weakly coupled cortical neuronal network. In all cases we find that the shotgun sampling succeeds in recovering the complete neuronal connectivity matrix from partial activity observations.

In Section 3.1, we prove important Theorem 1 that sets the basis for the subsequent investigation of the correctness of the shotgun connectivity estimation in different neuronal population activity models. This theorem also established important results about the design of possible shotgun-type neuronal connectivity imaging experiments. Theorem 1 establishes that any sparse neuronal activity imaging that covers a particular set of marginal “identifying” neuronal activity distributions is sufficient for the recovery of the complete connectivity matrix. In the linear model and the exponential-GLM of neuronal activity such sets of marginal neuronal activity distributions can be explicitly shown to consist of all time-shifted and same-time neuronal activity pairs, such as 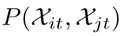 and 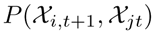. It is not important how such set of neuronal activity distributions was covered, what is important for the identifiability of full connectivity matrix is that such set is completely obtained. These results open new possibilities for investigating alternative shotgun-type neuronal population activity sampling organizations optimized for different experiment parameters.

In Section 3.2, we propose a particularly advantageous such alternative organization consisting of block-wise sequential scanning of a neuronal population’s activity. In this approach, the neurons are observed in two contiguous input and output blocks, sequentially, with all possible combinations of input and output blocks imaged during the experiment. This sampling organization guarantees the coverage of all same-time and time-shifted neuronal activity pairs and furthermore has the advantage of allowing straightforward implementations using existing fluorescent microscopy tools as well as simpler associated numerical connectivity estimation problem. At the same time, truly random sampling of the neuronal activity in a large neuronal population proposed in (Turaga et al., 2013; Keshri et al., 2013) is significantly more challenging, both from the experimental implementation stand-point and the solution of associated estimation problem. Thus proposed “block-wise round-robin” sampling strategy is simple conceptually and can be straightforwardly implemented experimentally, which hopefully will allow it to be realized in experimental designs by researchers in the near future.

Another particularly important issue raised by this study in Sections 3.3 and 3.4 is that of the scalability of the numerical shotgun connectivity estimation. We demonstrated here the shotgun connectivity estimation in simulated neuronal networks of up to *N* = 50 neurons. In practice, the reconstructions of real neuronal circuits will require solving this estimation problem for thousands and even millions of neurons. The SMC EM procedure described here is efficient, requiring *O*(*N*^2^*M*^2^*T*) time for the E-step and *O*(*N*^2^*MT*) time for the M-step, as well as parallelizable, making the solution of the above problems possible using the high-performance computing infrastructures currently existing in the world. Faster sampling schemes for hidden neuronal population’s activity may be proposed, for example, by using the fast Metropolis-Hastings algorithm described in (Mishchenko and Paninski, 2011), reducing the cost of the E-step to *O*(*N*^2^ *MT*).

At the same time, in this work the requirement to store up to *M* examples of the entire hidden neuronal population’s activity was the most significant burden on the numerical computation. For example, for *N* = 10^4^ neurons, *M* = 100 EM samples, and *T* = 10^4^ seconds of neuronal activity data recorded at 100 Hz, the required sample of the hidden neuronal activities constitutes and requires storing in the computer memory of staggering 10^12^ neuronal activity states. While this problem can be solved by partitioning over the nodes of a supercomputing infrastructure, research of alternative approaches for reducing the memory footprint of the shotgun connectivity estimation appears to be of relevance.

Among possible solutions is the use of alternative sampling schemes such as the block-wise round-robin strategy above, where the connectivity matrix estimation can be solved as a sequence of smaller problems, in which only the part of the experiment corresponding to a given input and output blocks is considered. While not affecting *N* and *M*, this reduces *T* to at most such containing the given input and output neuronal blocks. Another alternative is to make use of the sufficient statistics of the connectivity estimation problem in a given neuronal activity model. This would allow using the smaller sufficient statistics in the place of large hidden neuronal population activity samples. Finally, one other approach can consist in using approximate models for the combined inputs from hidden neuronal populations. For example, one can model the inputs into observed neurons from hidden population using a multivariate Gaussian distribution constructed using the current estimate of the connectivity matrix in the hidden and the observed neuronal population sectors.

Several issues remain open in this work and require future investigation. Among these is the issue of the identifying sets of marginal neuronal activity distributions for given neuronal population activity models. We proposed here, based on the degrees-of-freedom counting, that the sets of all pair-wise input-output neuronal activity distributions would suffice to uniquely constrain any network model of neuronal population activity parametrized by a single connectivity matrix 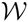 of *N*^2^ weights, where *N* is the number of neurons in the model. How can this conjecture be established rigorously? What are the identifying sets of the generalized linear model of neuronal activity defined by Eqs. (40-41) for general *f*(.)? What are the sufficient statistics of different network models of neuronal activity, that can be used in the connectivity matrix estimation? Do time-shifted correlation matrices provide the sufficient statistics for network models of neuronal activity?

Another set of questions is related to the design of the shotgun-like neuronal activity sampling strategies. Does the block-wise round-robin strategy offer the same speed of accumulating the information about different input-output neuronal pairs as the random sampling in the original shotgun proposal? What are other strategies that allow uniformly collecting pairwise inputoutput neuronal activity measurements? What are their important properties? Are all identifying sets equivalent from the point of view of accruing information about neuronal connectivity or some sets are more advantageous than the others? We hope that this work will stimulate further discussion regarding these questions.

## Acknowledgements

The author acknowledges the support via the TUBITAK ARDEB 1001 research grant number 113E611 (Ankara, Turkey), Toros University BAP grant number TUBAP135001 (Mersin, Turkey), and Bilim Akademisi—The Science Academy’s young investigator award grant under the BAGEP program (Istanbul, Turkey). The author wants to thank Liam Paninski for the key discussion leading to the development of this work, and Daniel Soudry for valuable comments on the manuscript’s draft. The author is also thankful to the anonymous reviewers, whose comments allowed the author to significantly improve the manuscript.

## A Sequential Monte Carlo Expectation Maximization algorithm for numerical solution of the shotgun connectivity estimation problem

The EM algorithm (Dempster et al., 1977) is the standard method of statistical inference in the presence of missing data. Briefly, the EM algorithm produces at least a locally maximum likelihood estimate of the parameters of a model *P*(*X, Y*|*θ*) given a set of observations *X* with the data *Y* missing, 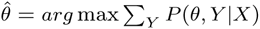. The EM algorithm produces a sequence of parameter estimates 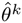 by iteratively maximizing the functions 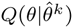,

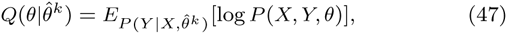
where 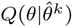 at each step is calculated by constructing *M* samples of the unavailable data *Y* from 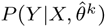 and using the following average,

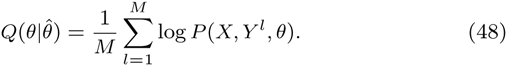

In the case of the shotgun sampling, the sampling step of the EM algorithm can be implemented using the forward-backward algorithm (Rabiner, 1989) and the sequential Monte-Carlo method also known as the Particle Filtering (Godsill et al., 2001). In this case, the distribution of the hidden neuronal activities at every observation is modeled by a sample of M hidden neurons’ activity configurations, 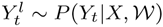, each referred to as a “particle”.

In order to produce this sample, it is advantageous to reformulate the sampling problem 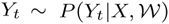 in a more convenient way as applying to drawing a sample of the complete neuronal activity configurations 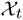 in such a way that the activity of the parts of the neuronal population observed at time *t* match the available neuronal activity data 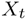. In this sense, we view the activity of the entire neuronal population 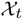 as “hidden” and the mapping of 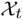 onto the subset of observed neurons, 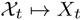, as the observation. In this form, the problem becomes that of sampling the sequence of the hidden states 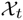 from a Hidden Markov Model with the observations *X_t_*. This problem now can be efficiently solved using the standard forward-backward algorithm.

Forward-backward algorithm consists of two passes. In the first forward pass, a sequence of samples of hidden neural activity states is produced according to 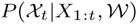, where *X*_1:_*_t_* refers to the collection of all observed neuronal activities up to and including the time *t*. Each sample in this sequence contains *M* examples of the complete neuronal population activity, 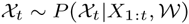, while the entire sequencecontains *T* such samples *t* = 1 … *T*, where *T* is the number of the observations, 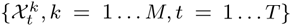.

Forward pass samples can be constructed iteratively by drawing the first sample 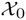 from the prior distribution 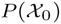, and then constructing each next sample according to,

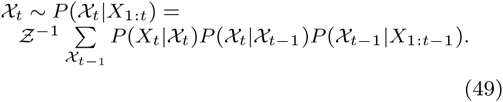

Here *Ƶ* is a normalization constant to be calculated below and we stopped writing parameter 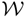 in the probability densities for brevity.

According to Eq. (49), the forward pass step at each *t* can be realized by taking the previous sample’s particles 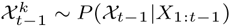 and ”moving” them according to the transition probabilities

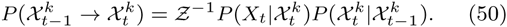

Eq. (50) can be simplified by noting that 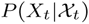 has the effect of only restricting the moves 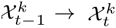 to such that make the activity patterns of the neurons observed in 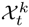 match the available observation *X_t_*,

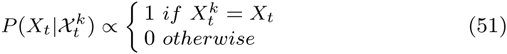

By using this and taking advantage of the factorization of the probabilities 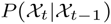 over individual neurons *i*, 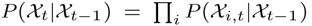, we can obtain the normalization constant *Ƶ* explicitly as,

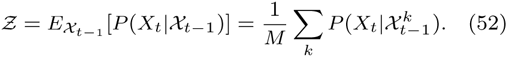

With this simplification, we arrive at the final forward step algorithm as follows:

### Forward Step

1. Select one 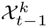 from the previous *t* − 1 sample 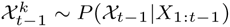 with the probability

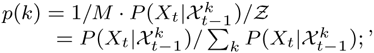
where

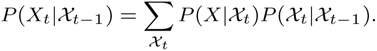
2. Set in 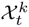 the activity of the neurons *i* observed in observation *t* as 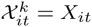
3. Set in 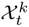 the activity of the neurons *i*ʹ not observed in observation *t* as 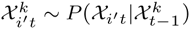.

In the backward pass, the samples 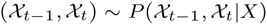 need to be constructed for each *t* conditional on the all observations *X* = {*X_t_, t* = 1 … *T*}. These samples can be constructed using the following relationship that we adopt here from (Paninski et al., 2010),

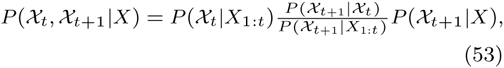
where 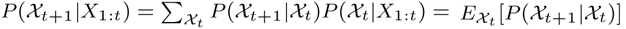, the average being over the forward pass sample 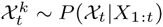.

According to Eq. (53), the backward step can be constructed by first combining into pairs the forward pass samples *t*, 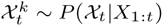, and the backward pass samples *t* + 1, 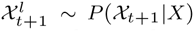, and then weighing these with the weights 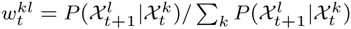. Evidently, thus formed pairs 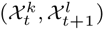 are distributed according to 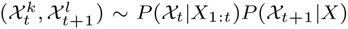, and the expectation value of any functional 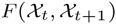 over 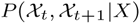 can be calculated by using such pairs as 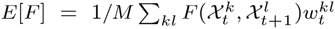. In addition, 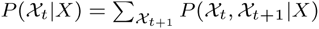 and the next backward pass sample for observation *t*, 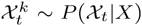, can be constructed by drawing with replacement 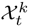 from 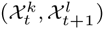 with probabilities 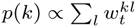.

Thus, we arrive at the final backward step algorithm as follows:

### Backward Step

1. Form *M*^2^ pairs 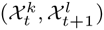 for each available forward pass sample 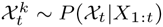 and backward pass sample 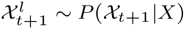;
2. Calculate the weights 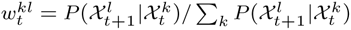
3. As the next backward pass sample 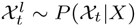 select with replacement 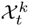 from the pairs 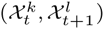 with the probabilities 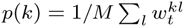;
4. The expectations values of a functional 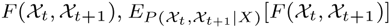, are given by 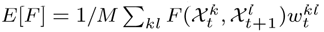.

In the optimization step of the EM algorithm we maximize with respect to 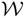 the following function,

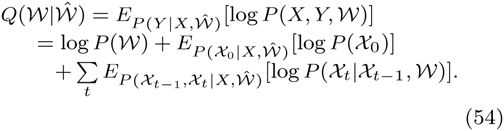

In order to calculate 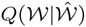 it is sufficient to know the samples 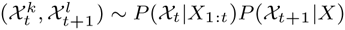 and the weights 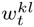. Moreover, 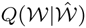 can be split into a sum over the rows of the matrix 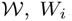, *W_i_*, as 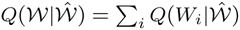, with 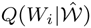 given by

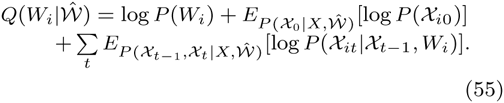

Thus, the optimization of Eq. (54) can be solved for each row *i* independently, reducing the complexity of the problem from quadratic in the number of neurons *N* to linear. Finally, inhomogeneous Poisson point-process models of neuronal activity with log-concave rate functions result in the problems 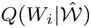 that are convex, which allows their efficient numerical optimization for very large *N* using the standard gradient descent methods (Paninski, 2004).

## B The calculation of posterior log-likelihood for the linear neuronal activity model with input neuronal populations.

In this appendix we calculate the marginal likelihood

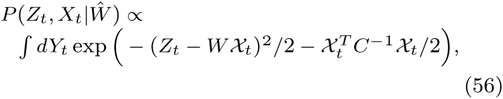
of the model (3), where the input neuronal activities are distributed according to a correlated Gaussian distribution with covariance matrix *C*,

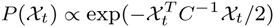
and the integration is performed over the part of 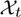, 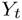, that is not observed during observation *t*. The part of 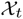 observed during observation *t*, respectively, is held fixed at *X_t_*. 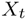 is a single row-vector from the full connectivity matrix 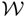 corresponding to the input connection weights of one “output” neuron.

The calculation of Eq. (56) can be simplified if we represent the integral in an invariant form by introducing *δ*-functions, which will restrict the integration over 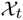 to the hyperplane of the observed neuronal activities *X_t_*, namely,

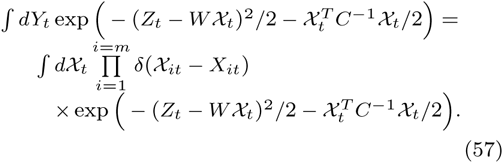

Here *m* is the number of the observed neuronal inputs and w.l.o.g. we assumed that the observed inputs *X_t_* comprise the first *m* elements of 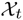. We now replace the *δ*-functions in Eq. (57) with their Fourier representation, 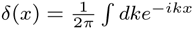, yielding

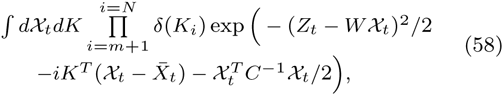
where 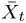 is a full-size column-vector of neuronal inputs, with the first *m* elements equal to *X_t_* and the rest of the elements zero (these do not affect the integral since *K_i_* = 0 for *i* > *m*). In Eq. (58), the integral over 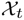 now can be taken explicitly as a Gaussian, resulting in

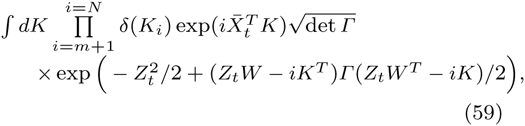
where the matrix *Γ* is identified from the part of the argument of the exponential in Eq. (58) quadratic in 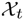, *Γ*^−1^ = (*C*^−1^ + *W^T^W*). We expand the second term under the exponential in Eq. (59) as

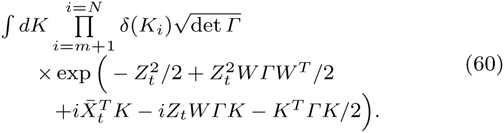

The *δ*-functions in Eq. (60) can be used subsequently to restrict the integration over *K* to only such values where *K_i_* = 0 for all *m* < *i* ≤ *N*. Thus, we rewrite this integration as

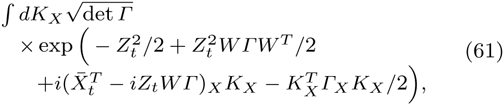
where the subscript *X* means restriction to the first m elements, as contained in the observed set of neuronal inputs *X_t_*. Thus obtained integration over *K_X_* is again Gaussian, and so we can perform it explicitly producing

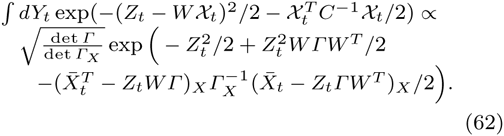

A simple check is in order now. By assuming *C = I* (uncorrelated inputs), we obtain by repeatedly using Woodbury lemma,

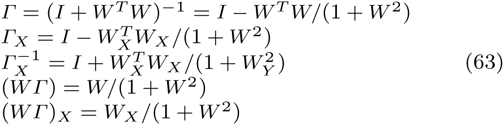
where *W_X_* and *W_Y_* are the restrictions of *W* to the subsets of the neuronal inputs *X_t_* and *Y_t_*, respectively, and *W*^2^ = *WW^T^*. For det *Γ* and det *Γ_X_*, we then obtain det *Γ* = (1 + *W*^2^)^−1^ and det 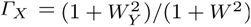, therefore,

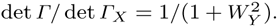

Similarly, we calculate

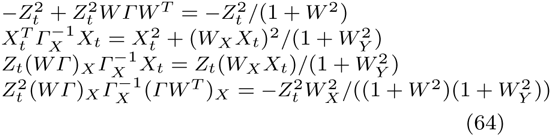

Combining all of Eqs. (64), we obtain for Eq. (61) and the case *C = I*,

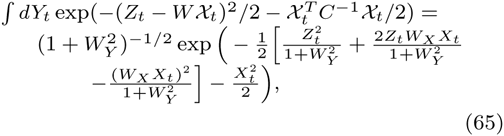
which is the same as Eq. (??) in the main text.

In the case of general *C*, we calculate similarly by repeatedly using Woodbury lemma

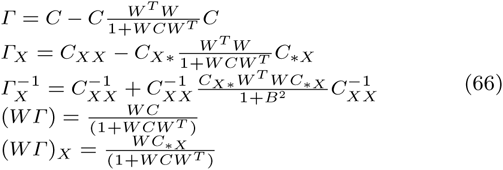
where

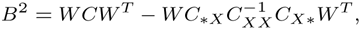
and *C_XX_* is the square block of the full covariance matrix *C* corresponding to the observed inputs *X_t_*, while 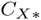 and 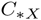 are the rectangular blocks of the full covariance matrix containing all the rows or the columns corresponding to the observed inputs *X_t_*. Then, for the determinant factor we obtain,

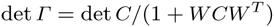
and

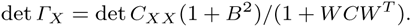

Consequently,

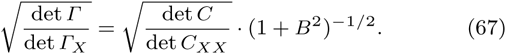

By considering

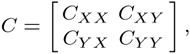
we can also reorder the quantity *B*^2^ as

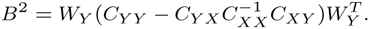

We see that *B*^2^ plays the role here of 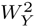 in Eq. (65). We further proceed to simplify all the terms under the exponent in Eq. (62) using Eqs. (66). This calculation is highly tedious, however, its result is obtained in the form,

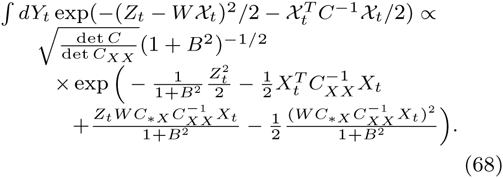

As a consistency check, we can verify that Eq. (68) reduces to Eq. (65) when *C = I*. Finally, we rewrite Eq. (68) more concisely as

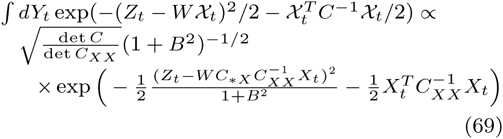

## Footnotes

1 In this respect, note that the pairwise same-time distributions 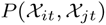 in general may not be sufficient to identify 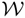 since these are symmetric and, in the worst case scenario, only provide *N*(*N* + 1)/2 independent constraints on 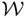. In that sense, the time-shifted distributions 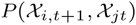 are guaranteed to provide *N*^2^ different constraints. One can also look at the distributions of triples 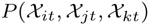 from 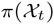. Alternatively, if the distributions 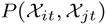 contain different statistics (such as mean and variance) that essentially independently relate to 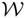, then measuring 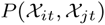 may also prove sufficient for identifying 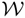.

